# Functional architecture of the pancreatic islets reveals first responder cells which drive the first-phase [Ca^2+^] response

**DOI:** 10.1101/2020.12.22.424082

**Authors:** Vira Kravets, JaeAnn M. Dwulet, Wolfgang E. Schleicher, David J. Hodson, Anna M. Davis, Laura Pyle, Robert A. Piscopio, Maura Sticco-Ivins, Richard K.P. Benninger

## Abstract

Insulin-secreting β-cells are functionally heterogeneous. Whether there exist cells driving the first-phase calcium response in individual islets, has not been examined. We examine ‘first responder’ cells, defined by the earliest [Ca^2+^] response during first-phase [Ca^2+^] elevation, distinct from previously identified “hub” and “leader” cells. First responder cells showed characteristics of high membrane excitability and lower electrical coupling to their neighbors. The first-phase response time of β-cells in the islet was spatially organized, dependent on the cell’s distance to the first responder cell, and consistent over time up to ∼24 h. When first responder cells were laser ablated, the first-phase [Ca^2+^] was slowed down, diminished, and discoordinated compared to random cell ablation. Cells that were next earliest to respond often took over the role of the first responder upon ablation. In summary, we discover and characterize a distinct first responder β-cell state, critical for the islet first-phase response to glucose.

## Introduction

Diabetes Mellitus is a disease characterized by high blood glucose, caused by insufficient secretion of insulin. β-cells within pancreatic islets of Langerhans secrete insulin and are compromised in diabetes. Early work showed that in mechanically-dispersed islets, single β-cells are heterogeneous in the level of insulin release [1]. More recent studies have discovered markers that separate β-cells into distinct populations with differing functional properties. This includes markers that subdivides proliferative-competent β-cells from mature β-cells [2]; subdivides β-cells with different levels of insulin gene expression, granularity and secretion [3]; or subdivides β-cells that have differing responsiveness to insulin secretagogues [4]. Furthermore, single-cell high-throughput approaches such as single cell RNA sequencing (scRNAseq) or mass cytometry separate distinct β-cell populations [5, 6]. However, the role of the heterogeneity in the function of the islet is poorly understood.

β-cells are excitable and show elevated electrical activity in response to glucose. Following metabolism of glucose, ATP-sensitive potassium channels (K_ATP_) close, depolarizing the membrane. This membrane depolarization opens voltage-gated calcium channels, elevating intra-cellular free Ca^2+^ activity ([Ca^2+^]) and triggering insulin release. β-cells are electrically coupled to neighboring β-cells via Connexin36 (Cx36) gap junction channels [7–11], which enables the direct exchange of cations between β-cells [12]. Under low glucose conditions, gap junction channels transmit hyperpolarizing currents that suppress islet electrical activity and insulin release [13–15]. Under elevated glucose gap junction channels coordinate the oscillatory dynamics of islet electrical activity, thereby enhancing first-phase insulin and pulsatile second-phase insulin, and glucose tolerance [11, 16].

Gap junction coupling is non-uniform throughout the islet [17]. β-cells are also heterogeneous in glucose metabolism and excitability [18]. As a result, at low glucose some β-cells are suppressed more than others by hyperpolarizing currents transmitted from neighboring cells [19]. Conversely at elevated glucose, some β-cells are recruited and/or coordinated more than others by depolarizing currents transmitted from neighboring cells. As such, the response of each β-cell within the islet to glucose is different, reflecting both its intrinsic heterogeneity and its context within the islet. Several studies have sought to identify and characterize functional β-cell states within the islet based on the [Ca^2+^] response under glucose stimulation, together with the use of optogenetic based constructs and laser ablation. For example, β-cells that show significantly increased connectivity, termed ‘hubs’ or ‘hub cells’ [20, 21], disproportionately suppressed islet [Ca^2+^] following targeted hyper-polarization via optogenetic stimulation. Conversely a population of β-cells disproportionately activated islet [Ca^2+^] following targeted depolarization via optogenetic stimulation [22]. Furthermore, cells that show [Ca^2+^] oscillations that precede the rest of the islet, termed ‘leader’ cells or ‘wave-origin’ [22–24], have also been suggested to drive the oscillatory dynamics of [Ca^2+^].

Both in rodents and in humans, first and second phases of insulin secretion have been distinguished [25, 26]. Glucose-stimulated [Ca^2+^] influx into the cell is necessary for both first and second phases of insulin secretion [27]. β-cell [Ca^2+^] is also bi-phasic [28] and is correlated with insulin secretion dynamics [29]. The first-phase of [Ca^2+^] response following low-to-high glucose is *transitional*, where cells with intrinsic differences in metabolic activity or other properties may be responding differently. The second-phase [Ca^2+^] response at high glucose is *steady-state* where the majority of cells are equally likely to fire [14], but with [Ca^2+^] oscillations showing differing amplitudes, temporal delay (phase lag), and oscillation frequency [30, 31]. While different β-cell subpopulations have been examined during this second-phase [Ca^2+^] response, the role of functional β cell states during the first-phase [Ca^2+^] has not been examined.

Here we identify functional β-cell state based on cell dynamics during the first-phase [Ca^2+^] response to glucose, and test whether they disproportionately affect islet function. We address what mechanisms underlie their properties and disproportionate effect on islet function and ask whether they overlap with other identified β-cell states.

## Results

### First responder cells are distinct from β-cell functional states associated with second-phase [Ca^2+^]

We first sought to identify β**-**cells that may exert control over the first-phase [Ca^2+^] response, and test whether they differ from cells reported to control the second-phase [Ca^2+^] response. [Ca^2+^] dynamics were recorded using confocal microscopy in the two-dimensional islet plane (Fig.1 A) continuously before and after glucose stimulation in islets isolated from Mip-Cre^ER^; Rosa-Stop-Lox-Stop-GCamP6s mice (β-GCamP6s) that show β-cell specific GCamP6s expression following tamoxifen-induced CreER-mediated recombination. We quantified the response time to glucose for each cell using initial [Ca^2+^] elevation (Fig.1 B) and defined the ‘*first responder*’ cells as 10% of cells in the islet with fastest response to glucose elevation, *T_resp_* (Fig.1 C and methods). The ‘*last responder*’ cells were defined as 10% of islet cells with latest response. The *T_resp_* distribution varied between the islets. The fastest 25% percentile preceded the islet-median response by ∼10 seconds, whereas the slowest 25% percentile lagged the islet-median by about the same amount of time (see cumulative *T_resp_* distribution for multiple islets in Fig.S1 A). First responder cells were located closer to the islet periphery (Fig.S1 B). Islets from the β-GCamP6s mice demonstrated a bi-phasic [Ca^2+^] response to glucose elevation, with the first phase characterized by a steep [Ca^2+^] rise and subsequent plateau (Fig.1 D red rectangle), and the second, oscillatory phase characterized by [Ca^2+^] wave (Fig.1 D blue rectangle). Some islets didn’t have a robust [Ca^2+^] wave (see methods).

**Figure 1.**
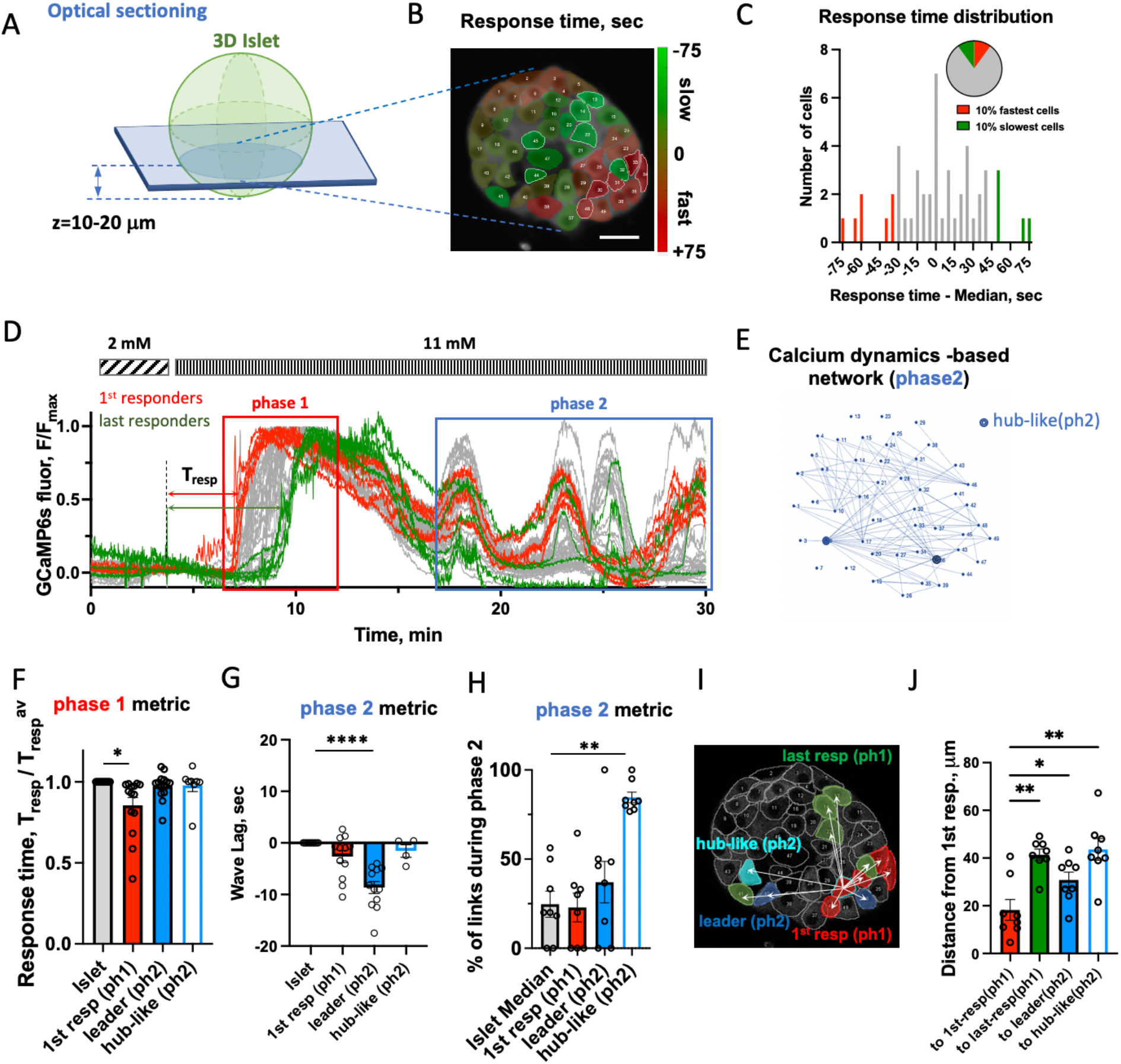
Identification of first responder cells. **A)** Schematic of where data was collected. **B)** Representative image of 2D islet plane with false-color map of response time to glucose elevation. **C)** Representative distribution of response time, Tresp, for the cells shown in (B). Pie chart describes definition of 1^st^ (last) responder cells as 10% of all cells in the islet 2D plane which have fastest (slowest) Tresp. **D)** Representative time-course of [Ca^2+^] dynamics within an islet shown in (B) following glucose elevation from 2 to 11 mM. In red rectangle: first-phase of the [Ca^2+^] response, in blue – second phase. Red curves correspond to first responder (1^st^-resp) cells, and green – to last responder (last-resp) cells. **E)** Link map based on the functional network analysis performed over the second phase dynamics in the islet shown in (B and E). Hub-like (ph2) cells are shown as blue dots. **F)** Phase1 [Ca^2+^] dynamics metric: response time following glucose elevation. Comparison of β-cells leading phase1 (1^st^ responder ph1) vs phase2 (leader ph2), vs hub-like (ph2) cells collected from 8-12 islets, 6-7 mice. **G)** Phase2 [Ca^2+^] dynamics metric: lag of the cell’s [Ca^2+^] wave with respect to the islet-average wave. Comparison of β-cells leading phase1 vs phase2 vs hub-like (ph2) cells (n=4-12 islets from 4-7 mice). **H)** Percent of functional links normalized to max number of links, identified during second phase of [Ca^2+^] dynamics (n=8 islets from 6 mice). **I)** Example of measurement of the distance from one of the 5 first responder cells in this islet to other cells. **J)** Quantification for multiple islets (n=8 islets from 6 mice), where each dot represents average distance from all first responder cells to other cell states per islet. Statistical analysis in F, G utilized one sample t test, H, J: utilized One-way ANOVA (with multiple comparison post-hoc test) where **** represents p<0.0001, *** p<0.0002, ** p<0.0021, * p<0.0332 for comparison of the islet-average value vs all other groups. Functional network analysis was performed via binarization and co-activity matrix analysis, as described [20]. See Figure 1 - Source Data for values used in each graph.

We next examined the overlap between properties of cells defined by the first-phase and second-phase [Ca^2+^] dynamics. During the oscillatory second phase, highly connected “hub cells” have been implicated in maintaining [Ca^2+^] elevations. To assess whether first responder cells overlap with these “hub” cells, we identified “hub-like (phase2)” cells via network analysis as performed previously [20] (Fig.1 E). ‘*Leader*’ cells, which have previously been defined and characterized [22, 24], were defined as 10% of all cells with highest *negative* phase lag, which corresponds to showing an earlier [Ca^2+^] oscillation (Fig.S1 C, D).

The response times of leader and hub-like(ph2) cells were no different from the islet average (Fig. 1F), *i.e.,* they did not lead the first-phase [Ca^2+^] response to glucose. While the first responder cells had significantly different *T_resp_* compared to the islet-average cell (p=0.0105). During the second-phase [Ca^2+^] dynamics, the *phase lag* of the [Ca^2+^] wave of the hub-like(ph2) cells was no different from the islet average (Fig1 G), *i.e.,* they did not lead the second-phase [Ca^2+^] response to glucose. Interestingly, the first responder and leader cells had phase lag which was substantially different from that of islet-average (p=0.0476 and p<0.0001 correspondingly), indicating that they both lead second-phase [Ca^2+^] dynamics. However, further analysis showed significant difference between these two cell states: leader cells had much earlier calcium wave (greater negative phase lag) compared to the first responder cells (p=0.0109). Thus, based on the first- and second-phase [Ca^2+^] dynamic metrics, first responder cells were distinct from the leader cells.

Comparing coordination of [Ca^2+^] wave during the second-phase [Ca^2+^] response to glucose, we found that first responder and leader cells were not different from the islet-average (Fig.1 H), unlike hub-like(ph2) cells which showed a significantly higher number of links compared to the islet average (p=0.0018). Thus, first responder cells are distinct from previously defined hub cells. Extended analysis of the features of the [Ca^2+^] dynamics (including network analysis revealing hub-like cells in the first-phase [Ca^2+^]) is presented in Fig.S1 E-K.

We next looked at the spatial organization of the first-phase [Ca^2+^] response to glucose. Distances between different β-cells were measured in a z-plane ∼10-20 μm away from the surface of the islet’s beta-cell core. (Fig.1A). Distances were measured from each first responder cell in the islet and then averaged (see Fig.1 I, J and methods). First responder cells formed clusters of size <20μm (Fig.1 J red bar). The first responder clusters were significantly spatially separated from leader cells (p=0.0208) and from hub-like(ph2) cells (p=0.0056). On average first responder cells were located <30μm away from leader cells, and 45 μm from hub-like(ph2) cells (Fig.1 J). The response time of β-cells in the islet following glucose stimulation was generally dependent on their relative proximity to a first responder cell, where cells closer to first responder cells responded earlier (Fig.S2 A). A linear dependence of the response time vs proximity of the first responder was clear for some islets. For other islets location of the first responder cells was such that multiple response domains with a local first responder in the center were formed (examples are shown in Fig.S2 B). We defined the speed of the response propagation as the slopes of the distance to first responder vs the time of response (Fig.S2 A). This time over which the initial response to glucose response propagated across the islet was measured here for the first time, and it varied substantially from islet to islet between 0.16 and 3.3 μm/sec. Given different islets had substantially different rates of response to glucose we subsequently normalized the response time by islet.

### Clusters of first responders are spatially consistent

We reasoned that if cells respond randomly to glucose stimulation, then first responder cells identified during the initial glucose elevation will not be the same first responder cells during a repeated glucose elevation. Conversely if there exists a functional hierarchy of cells within the islet, then there should be consistency in the location of the first responder cells. We first stimulated islets to elevated (11 mM) glucose, lowered back to basal levels, and re-stimulated over the course of 1-2 h (Fig.2 A). The location of the first responder cluster in the islet remained consistent between the initial and repeated glucose elevation (Fig.2 B, C). Some initial first responder cells were first responders in the repeated glucose stimulation (35% of cases), whereas some surrendered their role either to the nearest neighbor cell (35%), or to the 2nd neighbor (18%), or to a more distant cell (12%).

**Figure 2.**
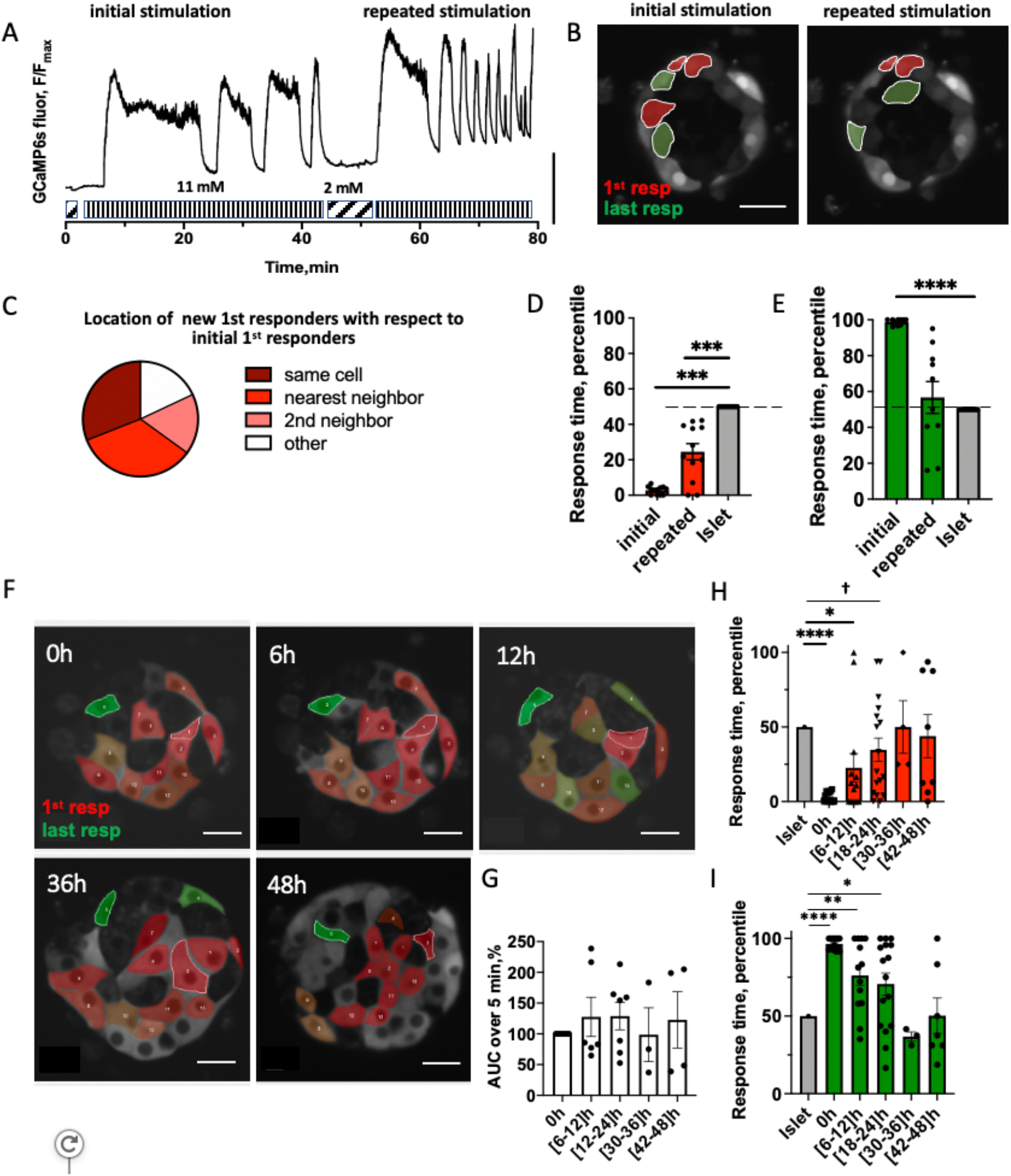
Consistency of first responder cells. A) Representative time-course of [Ca^2+^] dynamics within an islet under repeated glucose stimulation. Vertical scale bar represents 30% of the fluorescence intensity change. B) Representative image of the islet with the location of first responders (red) and last responders (green) during initial and repeated glucose elevation. C) Spatial consistency: Spatial location of new first responder cells (during repeated glucose elevation) relative to the old first responder cells (during original elevation) (n=10 islets from 4 mice). D) time of response of cells identified as first responders during the initial glucose elevation and upon repeated glucose elevation (n=10 islets from 4 mice). E) As in D for last responders. F) False-color map indicating [Ca^2+^] response time to glucose elevation during the first phase, recorded in the same region of the islet over 48 hours at 6 hour intervals. Scale bar indicates 20 μm. G) Area under the curve for [Ca^2+^] elevation over 0-5 min for the islet average, for each time-window indicated. Data is normalized to the [Ca^2+^] AUC at 0h. H) time evolution of the response time of the cells identified as first and I) As in H for last responders (n=7 islets from 3 mice). Statistical analysis in D, E utilized one-sample t-test (with the null hypothesis of initial or repeated difference from the islet being 0). H, I utilized LMEM. † in H indicates p=0.06. See Figure 2 - Source Data for values used in each graph and Statistical Analysis – Source data for LMEM and one sample t-test details.

We examined location of the first- and last responder cells, as well as leader cells in 3-dimensions (Fig. S3). Within three cell layers, each separated by 10 μm, the locations of each β-cell subpopulation were conserved, suggesting functional organization in 3-dimensions, as we observed in 2-dimensions.

### Temporal consistency of first responder cells in not rigid

We next sought to determine whether an early response time was a consistent feature of first responder cells. As above, we first stimulated islets to elevated (11 mM) glucose, lowered back to basal levels, and re-stimulated over the course of 1-2 h (Fig.2 A). The period of the observed slow oscillations was in the same range as previously reported [37], (T=246±89 sec, Fig.S4 A). No significant difference was observed in the frequency of the oscillations during the repeated stimulation (0.011±0.008 Hz vs 0.007±0.001 Hz, Fig.S4 B). Response times for all cells during the initial and repeated glucose stimulation are shown in Fig.S4 C for individual islets. When considering all cells within the islet, there was no significant correlation between the response time of a cell during the initial stimulation compared to the response time upon the repeated stimulation. However, first responder cells in the majority of islets still remained consistent, while last responder cells were not consistent (Fig.S4 C). We found that temporal consistency of *first responder* cells (Fig.2 D) was substantial, but not fully rigid. Upon re-stimulation, the initial first responders retained a response that was significantly earlier than the median islet response time: on average in the fastest 25% of the whole *T_resp_* distribution. In contrast, the last-responding cells lacked consistency upon repeated glucose elevation, with a response time close to the islet-average (Fig.2 E).

To test whether first responder cells are maintained over a longer time, we measured the response time upon elevated glucose within the same cell layer in the islet for 48 h at 6 h intervals. At each time point we defined the first responder and last responder cells (Fig.2 F). There was no significant difference in the total [Ca^2+^] influx in the islet at each time point (Fig.2 G). The first responder cells remained consistent during the first 12 hours, where they showed significantly earlier [Ca^2+^] response time than the islet average. However, at 18-24 h the first responder cells became less distinguishable from the islet average, and at >24 h, their response time was indistinguishable from the islet-average (Fig.2 H). A similar temporal pattern was observed for the last responder cells (Fig.2 I). Thus, not all β-cells responded randomly to glucose stimulation. Rather, a first responder β-cell state consistently led this response, but this was maintained only over a ∼24h time period.

### First responder cells show characteristics of more excitable cells

β-cells that have been previously identified to disproportionately recruit or maintain elevated [Ca^2+^] in neighboring cells have shown elevated excitability, such as arising from increased metabolic activity [22]. We next examined characteristics of the first responder cells that may allow them to show an earlier response time. We first tested whether first responder cells have differing regulation of resting membrane potential (for example, a lower K_ATP_ conductance or increased depolarizing current) that would increase the likelihood of earlier membrane depolarization in response to glucose elevation. We performed sequential glucose and glibenclamide stimulation, in the same manner as repeated glucose stimulation experiments. Following each stimulation, we identified first responder cells (Fig.3 A). Those cells which responded first to glucose, also showed a significantly lower than average response time to glibenclamide (Fig.3 B, C) (p=0.046). In contrast, those cells which responded last to glucose, did not show a response time different to the islet average under glibenclamide (Fig.3 D). When all cells in the islet plane were considered, we did not observe a significant correlation between the response time during glucose stimulation as compared to the response time upon glibenclamide stimulation. This suggested that while differing K_ATP_ conductance (or resting depolarizing current) may be one factor in defining the earlier response time for a *first responder cell*, other factors are also likely involved.

**Figure 3.**
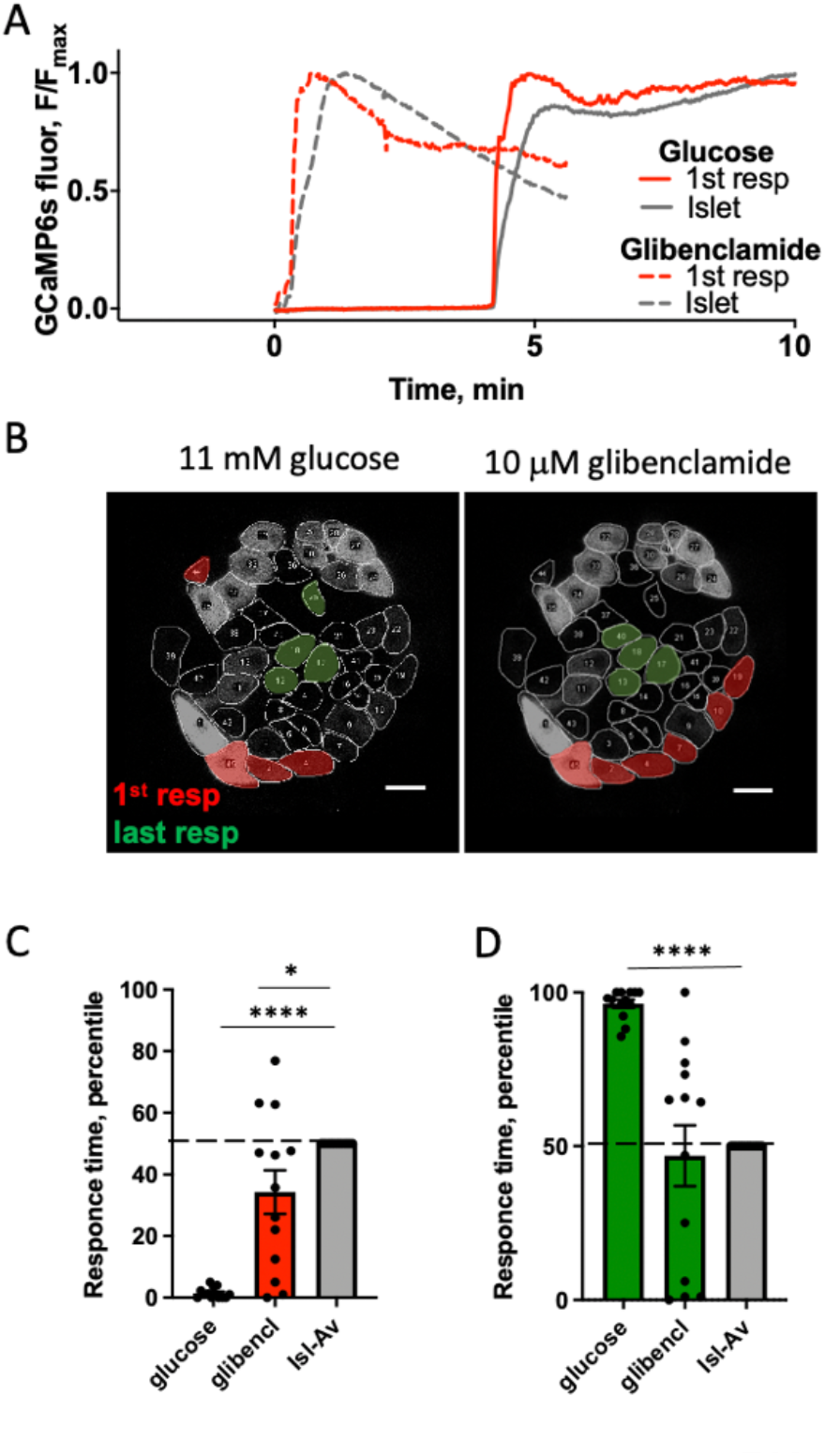
Responsiveness of first responder cells to secretagogues. A) Representative time-course of [Ca^2+^] dynamics within an islet under 11 mM glucose stimulation (dashed curve) and under 10 μM glibenclamide (solid curve) stimulation for first responder cell (red) and islet-average (grey). B) Representative image of the islet with the location of first responders (red) and last responders (green) during glucose and glibenclamide stimulation. Scale bar indicates 20 μm. C) time of response of cells identified as first responders during glucose stimulation and upon glibenclamide stimulation (n=13 islets from 7 mice). D) As in C for last responder cells. Statistical analysis in C, D utilized one-sample t-test (with the null hypothesis of glucose or glibenclamide difference from the islet being 0). See Figure 3 - Source Data for values used in each graph.

To discover other factors that may characterize first responder cells, we examined the [Ca^2+^] influx upon glucose stimulation (Fig.4 A). First responder cells demonstrated significantly higher influx of [Ca^2+^] during the first-phase [Ca^2+^] response (p=0.0072), while last responder cells had significantly lower [Ca^2+^] influx (p=0.453) (Fig.4 B, C). First responder cells did not have a greater than average NAD(P)H levels at either low (2 mM) or elevated (11 mM) glucose (Fig.4 D, E). Interestingly, following dye transfer kinetics via FRAP first responder cells showed lower than average gap junction permeability (p=0.0270) (Fig.4 F, G).

**Figure 4.**
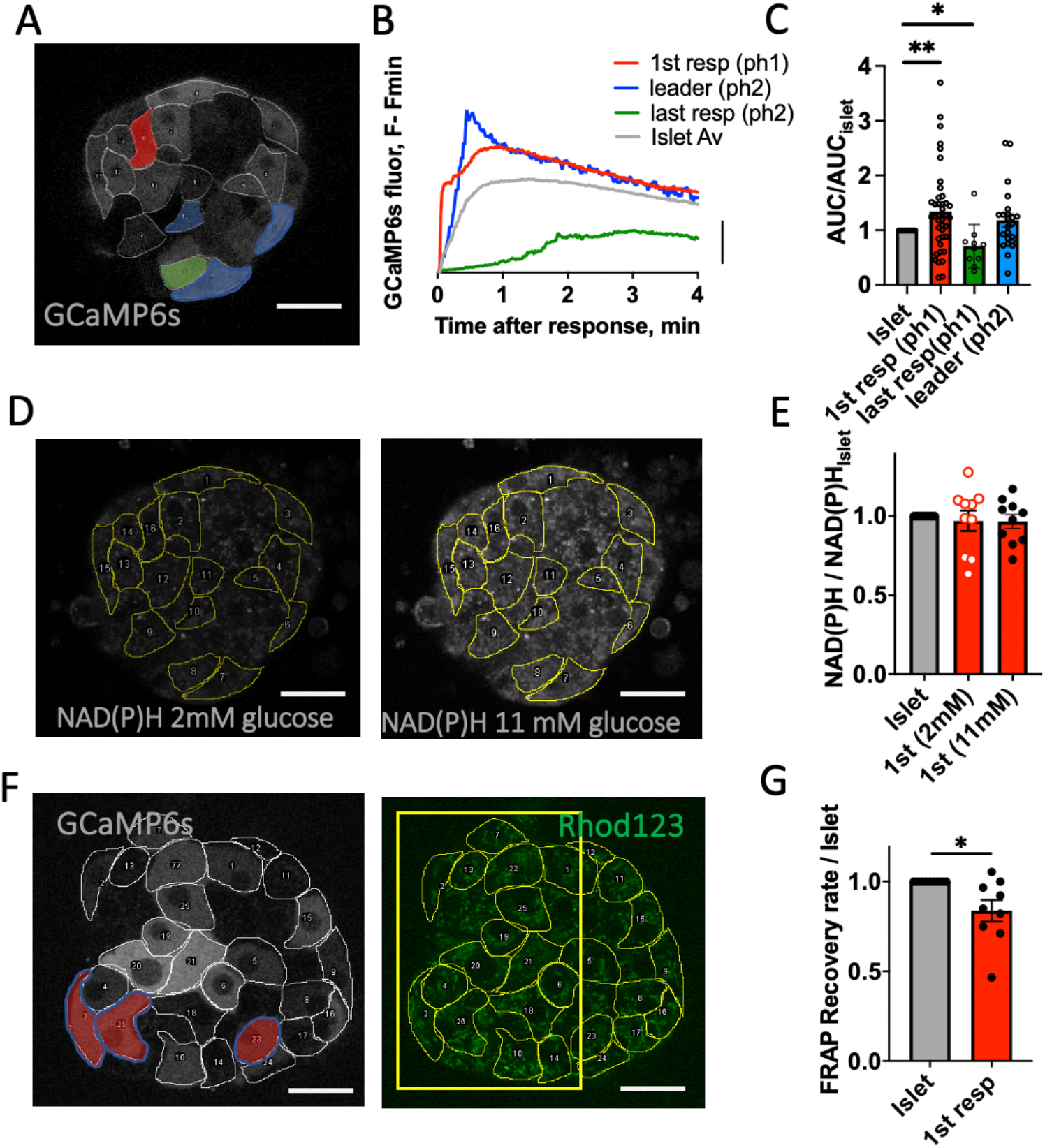
Characteristics of first responder cells. A) Representative image of the islet with first responders (red), leader (blue) and last responder (green) indicated. B) Time course of [Ca^2+^] elevation in each cell population, indicating Area under the curve. C) Area under the curve for each cell state during the first 3 min after [Ca^2+^] elevation upon elevated glucose (16 islets from 9 mice). D) Representative image of NAD(P)H autofluorescence in islet indicated in A, at low (2 mM) and high (11 mM) glucose. E) Mean NAD(P)H intensity in each identified subpopulation of β-cell, at low (2 mM) and high (11 mM) glucose, normalized with respect to the islet average (n=10 islets from 10 mice); F) Reprehensive GCamP6s fluorescence and Rhodamine123 fluorescence (for FRAP measurements) within the same islet layer. G) Mean Rhodamine123 fluorescence recovery rate as calculated during FRAP (n=7 islets from 5 mice). Scale bar indicates 20 μm. Statistical analysis in C utilized one-sample t-test where **** represents p<0.0001, *** p<0.0002, ** p<0.0021, * p<0.0332 for comparison of the islet-average value vs all other groups. In E, G utilized one-sample t-test (with the null hypothesis of 1^st^ responder difference from the islet being 0) where * p<0.05 indicated comparing the islet-average value vs first responder group. See Figure 4 - Source Data for values used in each graph.

### First responder cells drive first-phase [Ca^2+^] elevation

To test whether the hierarchy of cell responsiveness is functionally important, and specifically whether the earliest responding cells disproportionally drive the islet [Ca^2+^] response, we removed single β-cells from the islet via two-photon induced femtosecond laser ablation. Two-photon laser ablation allows for highly targeted removal of a cell without disrupting cells in close proximity [33]. As previously, we measured [Ca^2+^] dynamics upon elevation from 2 mM to 11 mM glucose; lowered glucose back to 2 mM, ablated one cell, and then repeated glucose elevation (Fig.5 A). Under each glucose elevation we identified first responder cells (Fig.5 B). Upon ablation of a control (non-first responder) cell the islet [Ca^2+^] response was relatively unchanged, with robust second-phase [Ca^2+^] oscillations (Fig.5 A, B). Upon ablation of a first responder cell robust second-phase [Ca^2+^] oscillations were also observed (Fig.5 C). And a new cell within the islet becomes the first responder cell (Fig.5 D).

**Figure 5.**
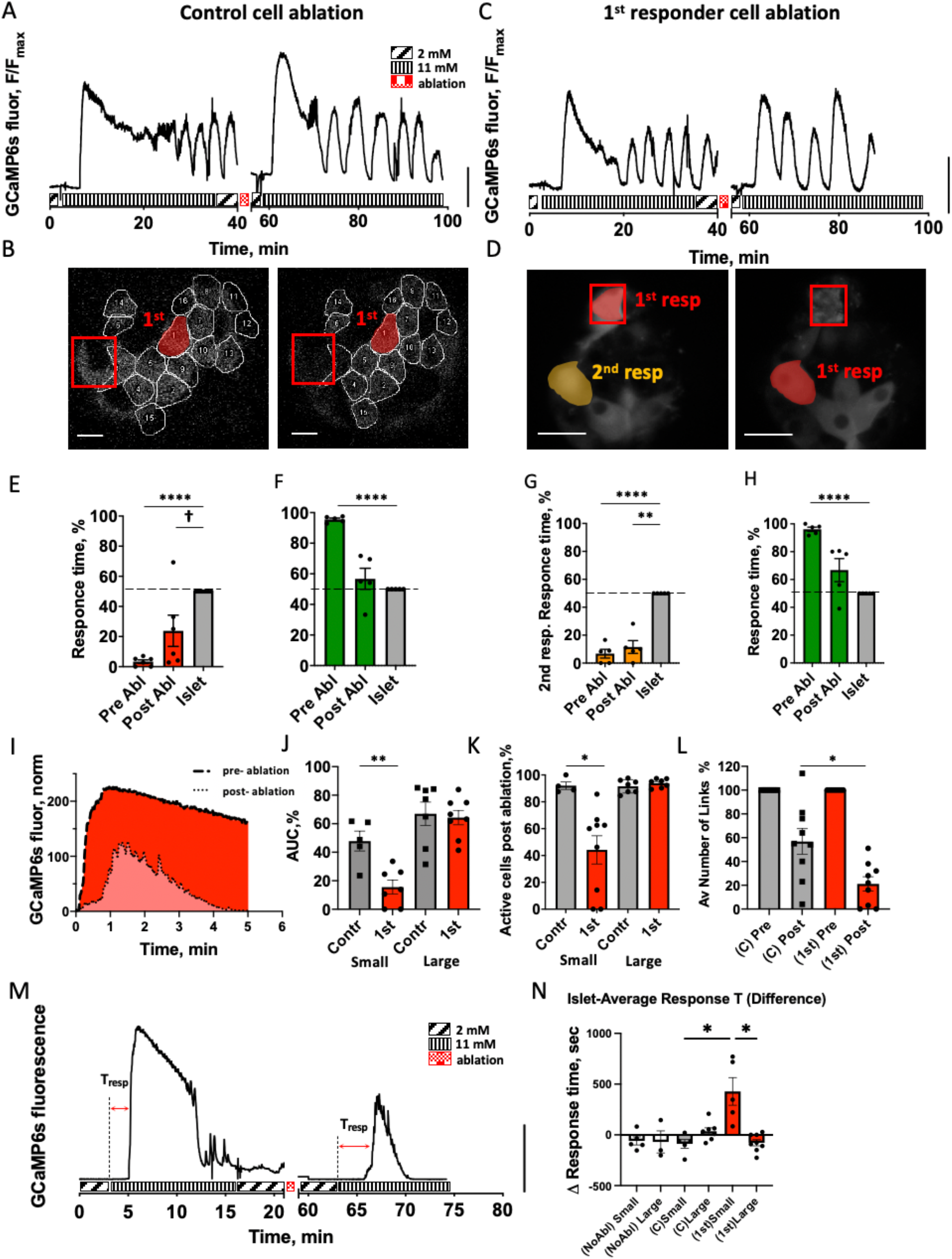
Role of first responder cells. **A)** Representative time-course of [Ca^2+^] dynamics within an islet pre- and post-ablation of a control cell (non-first responder cell). Vertical scale indicates 30% fluorescence intensity change. **B)** Representative image of the islet with the location of first responders (red) pre- and post-ablation of a control cell (red square). **C)** As in A for ablation of first responder cell. **D)** As in B for ablation of first responder cell (Red square), with second responder cell highlighted (orange). **E)** time of response of cells identified as first responders prior to control cell ablation and following control cell ablation (n=6 islets from 6 mice). **F)** As in E for last responders. **G)** Time of response of cells identified as second responders prior to first responder cell ablation and following first responder cell ablation (n=5 islets from 5 mice). **H)** As in G for last responders. **I)** Representative time course of Ca^2+^ elevation following glucose elevation pre- and post-ablation, with the Area-Under-the-Curve indicated. **J)** Area-Under-the-Curve for the first-phase [Ca^2+^] response post ablation of a control and first responder cells, for small (<89 μm diameter) and large (>89 μm diameter) islets. Data is normalized to the Area-Under-the-Curve prior to ablation. (27 islets from 17 mice) **K)** % of cells that show [Ca^2+^] elevation post ablation of a control and first responder cells, for small and large islets. Data is normalized to the Area-Under-the-Curve prior to ablation; **L)** Average number of functional links per islet during the first-phase of [Ca^2+^] response for a control and first responder cell ablations, for small and large islets combined. Data is normalized to the number of functional links prior to ablation (n=9 islets for each group). Functional network analysis was performed via Pearson-product algorithm, as described before [21]. **M)** Representative [Ca^2+^] pre- and post-first responder ablation in a small islet, Tresp indicates islet-average response time. **N)** Response time change in case of no ablation (white bars), control cell ablation (grey bars), or first responder cell ablation (red bars) (35 islets from 20 mice). 2 small islets that underwent first responder cell ablation did not respond at all following ablation and were excluded from analysis due to “infinite” response time post-ablation. Statistical analysis in E, F, G, H utilized one-sample t-test (with the null hypothesis of pre-or post-difference from the islet being 0) where **** represents p<0.0001, *** p<0.001, ** p<0.01, * p<0.05 indicated for comparison of the groups. † in (E) indicates significance of p=0.06. In I, J, K, L utilized ANOVA with multiple comparison Kruskal-Wallis test, where * p<0.0332 comparing the groups indicated. See Source Data – Figure 5 and Statistical analysis files for values used in each graph.

We examined the [Ca^2+^] response timing and [Ca^2+^] elevation in cells across the islet during the first-phase [Ca^2+^] response to glucose following ablation of either control (non-first responder) cell or first responder cell. Following ablation of the control cell, the response time of first responder cells remained below the islet-average (Fig.5 E, p=0.0022); whereas last responder cells were not different from the islet average (Fig.5 F). On average 53% of the original first responder cells remained as the earliest responding cells (Fig.S5 A center). These findings are very similar to those in the absence of any cell ablation (Fig.S5 A left). Following ablation of a first responder cell, the response time of the next earliest responder cells remained below the islet-average (Fig.5 H, p=0.0019); whereas last responder cells again were not different from the islet average (Fig.5 I). On average 40% of the original next earliest responding cells (‘second responder’ cells) become new earliest-responding cells (Fig.54 A right) and in another 20% of cases they remained as second responder cells. This suggests a hierarchy in the timing and β-cell responsiveness to glucose.

We next quantified the effect of a cell ablation on the whole islet first-phase [Ca^2+^] response. We first defined the [Ca^2+^] influx as the area under the curve for [Ca^2+^] over 5 min following elevation (Fig.5 K). We observed a size-dependence in area under the curve in both control and first responder ablation cell experiments (Fig.S5 B), hence we separated islets into small and large islets based on the median of the size distribution (Fig.S5 C). In large islets (>89 μm diameter) the islet-average [Ca^2+^] influx was not different between control or first responder cell ablation cases (Fig.5 J). In small islets (<89 μm diameter) while the islet-average [Ca^2+^] influx decreased following ablation of each cell type, the decrease was significantly greater following ablation of first responder cells. Consistent with these observations the proportion of cells that showed elevated first-phase [Ca^2+^] (i.e. within 5 min. of [Ca^2+^] elevation) was largely unchanged in large islets, with 93% of cells showing elevated [Ca^2+^] for ablation of both first responder or control cells (Fig.5 K). In small islets the proportion of cells that showed elevated first-phase [Ca^2+^] was largely unchanged following control cell ablation (91% of cells), but substantially decreased following first responder cell ablation (42% of cells). A similar trend was observed in terms of connectivity during the first-phase [Ca^2+^] elevation: the mean number of functional links within the islet was significantly less following first responder cell ablation compared to the control cell ablation (Fig.5 L). Finally, we measured the response time of the islet following ablation of a first-responder cell (Fig.5 M, N). We did not observe significant changes in large islets under ablation of a first responder cell, control cell or with no ablation. Similarly, we did not observe significant change in small islets under ablation of a control cell or with no ablation. However, following ablation of a first responder cell, the time for a [Ca^2+^] response of an islet significantly increased (Fig.5 M, N). Thus, in smaller islets, removal of a first responder cell via femtosecond laser ablation diminishes and delays the elevation of first-phase [Ca^2+^] as a result of there being fewer cells that respond both rapidly and in a coordinated fashion.

### Computer model description of first responder cell characteristics

To understand which characteristics of first responder cells are required for their action we utilized a previously published multi-cell model of islet β-cell electrophysiology [34]. This model incorporates heterogeneity in multiple metabolic, electrical and gap junctional characteristics. We simulated the [Ca^2+^] response upon elevated glucose (Fig.6 A) and observed significant variability in the time of [Ca^2+^] elevation (Fig.6 B). We defined first responders as the 10% of all cells with the fastest response times, as in experiments. We first examined the characteristics of these first responder cells in the model islet. The rate of glycolysis for model first responder cells was not significantly different from the islet-average (Fig.6 C), consistent with NAD(P)H measurements. The open channel K_ATP_ conductance (equivalent to channel number) for model first responder cells was significantly lower than the islet-average (Fig.6 D), in agreement with the earlier-than average response time under glibenclamide stimulation. The coupling conductance for model first responder cells was slightly, but significantly lower than the islet average (Fig6 E), consistent with gap junction permeability FRAP experiments. Model last responder cells showed no difference in glycolysis rate, elevated K_ATP_ conductance and reduced coupling conductance (Fig.6 C-E). Thus, model first responder cells are characterized by a combination of both lower electrical coupling and higher membrane excitability because of lower K_ATP_ conductance; whereas model last responder cells are characterized by a combination of both lower electrical coupling and lower membrane excitability because of greater K_ATP_ conductance.

**Figure 6.**
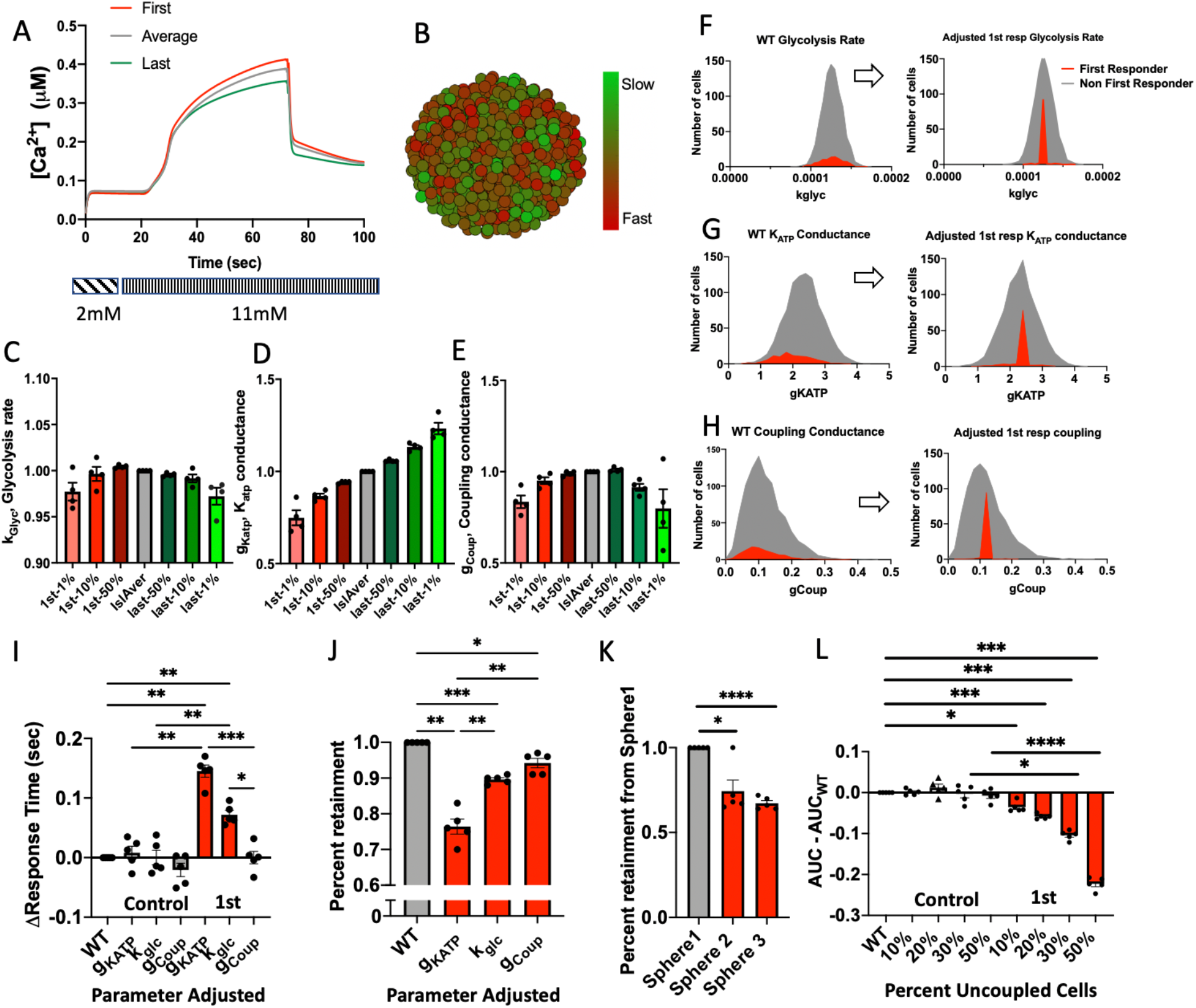
Modelling first responder cells. A) Representative time-course of [Ca^2+^] in a simulated islet, indicating the [Ca^2+^] dynamics characterizing the first responder (1^st^-resp) and last responder (last-resp) cells compared to the islet-average. B) False-color map indicating [Ca^2+^] response time to glucose elevation during the first phase. C) Mean parameter value of glycolysis rate (k_glyc_) averaged over the first (red) and last (green) indicated % of responding cells (n=4 seeds). Data is normalized with respect to the islet average parameter. D) As in C for K_ATP_ conductance (g_KATP_). E) As in C for coupling conductance (g_coup_). F) Distribution of the glycolysis rate (k_glyc_) in non-first responders (grey) and first responders (red) before (left) and after (right) parameter adjustment to set first responder parameter the same to the islet average value (n=5 seeds). G) As in F for adjustment of K_ATP_ conductance (g_KATP_). H) As in F for adjustment of coupling conductance (g_coup_). I) Change in response time upon adjustment of the glycolysis rate (k_glyc_), K_ATP_ conductance (g_KATP_) and coupling conductance (g_coup_) in first responder indicated in F-H, or in a random sub-set of cells. J) Percent of the original first responder cells that remain among the earliest responders after adjustment of adjustment of k_glyc_, g_KATP_, g_coup_ in first responder cells indicated in F-H. K) Percent of the original first responders that remain among the earliest responders after adjustment of the spatial position of the first responders in three separate distributions (spheres) (n=5 seeds). L) Decrease in the Area-Under-the-Curve (AUC) of the first phase of the Ca^2+^ response following de-coupling (removal) of a given % of the first responder or a random set of cells from the islet (n=5 seeds). Statistical analysis in I, J, K, L utilized one-way ANOVA (with Multiple comparison post-hoc test) where **** represents p<0.0001, *** p<0.001, ** p<0.01, * p<0.05 comparing he groups indicated. See Figure 5 - Source Data for values used in each graph.

To investigate the relative role of these functional characteristics and to define first responder cells, we simulated the [Ca^2+^] response following each of the above-mentioned parameters being *adjusted* to be equal to the corresponding islet-average parameter (Fig.6 F-H): little change in glycolysis rate, increase in K_ATP_ conductance or increase in coupling conductance. Thus, if a specific parameter is necessary for the function of the first responder, adjusting that parameter would be expected to significantly change the response time of that cell. Of the three parameters examined, re-adjusting the K_ATP_ conductance had the greatest impact on the first responder cell response time (Fig.6 I) and whether a first responder cell retained the earliest [Ca^2+^] elevation (Fig. 6J). In addition, rearrangement of the cell position within the islet, and thus the neighboring cells in contact with the first responder cell, significantly disrupted whether a first responder cell showed the earliest elevation in [Ca^2+^] (Fig. 6K). Thus, model first responder cells are defined by both increased excitability and position within the islet.

Finally, we tested whether first responder cells acting via gap junction coupling were sufficient to recruit neighboring cells. Following simulation of the islet, we removed either a random set of control cells or the earliest responding cells and re-simulated the islet [Ca^2+^] response. Consistent with experimental measurements, removal of first responder cells diminished the area under the curve for the islet-average [Ca^2+^] elevation, whereas removing a control set of cells had no impact on the islet-average [Ca^2+^] elevation (Fig.6 L). Importantly a decrease in [Ca^2+^] elevation was only observed for removal of greater than 10% of the earliest-responding cells; with greater reduction for removal of a higher % of cells. Thus, gap junction coupling is sufficient for a population of first responder cells to exert control over first phase [Ca^2+^] in the islet.

## Discussion

β-cells within the islet are functionally heterogeneous. While subpopulations of β-cells have been suggested to maintain coordinated oscillatory [Ca^2+^] and insulin release [35], these are associated with the second phase of the [Ca^2+^] response; a point at which cell-cell electrical communication is less important for *recruiting* [Ca^2+^] and the level of insulin release, but is critically important for *coordinating* pulsatile [Ca^2+^] and insulin secretion dynamics. Given the importance of cell-cell electrical communication in regulating first-phase insulin release [16], we examined whether there exists a subpopulation of β-cells associated with the first-phase calcium response. We discovered a functional cell state that we termed “first responder”. First responder cells lead the first-phase [Ca^2+^] response, and were distinct from previously identified functional subpopulations of β-cells. These cells were more excitable, critical for recruiting β-cells to elevate [Ca^2+^] immediately following glucose stimulation (Fig.7). We further demonstrated that this state of the β-cell is conserved over a ∼24 h time period.

**Figure 7.**
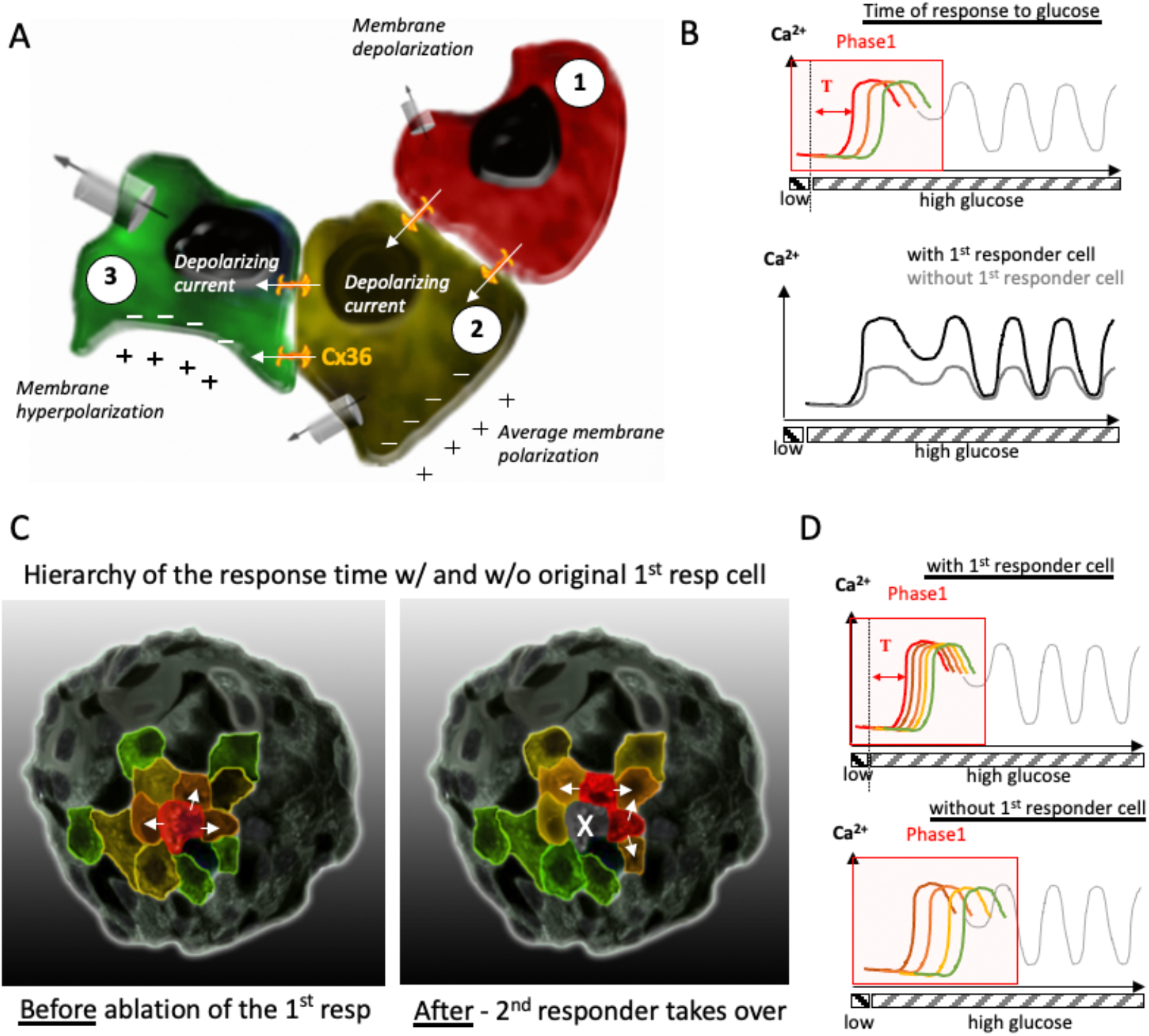
Schematic representation of the role of first responder cells in the islet. A) Cell1 - first responder cell (red) is more prone to membrane depolarization upon glucose stimulation. Cell 2 (yellow) is less depolarized than a first responder. The difference in the membrane potential between the cell 1 and 2 leads to depolarizing current to flow through the gap junctions into cell 2 triggering depolarization. Cell 2 subsequently depolarizes the less excitable cell 3 (green). B) Representation of the [Ca^2+^] response to glucose in cells 1,2,3 shown in (A), and the islet-average [Ca^2+^] response with (black) and without (grey) the first responder cell. C) Schematic of the time of response in the islet (red – faster response, green - slower) before and after the first responder ablation. Post-ablation, the cell with the second-earliest response time takes over the role of the first responder. D) Representation of the [Ca^2+^] coordination before and dis-coordination after the first responder cell ablation.

### First responder cells represent a distinct functional state of the β-cell

We defined first responder cells as those β-cells that lead the first-phase [Ca^2+^] response, more specifically the 10% of cells in the islet with the earliest response time. Importantly upon repeated stimulation, the same cells generally showed an earlier-than-average response over 1-2 hours (Fig.2), unlike last responder cells, which lacked any consistency. Only in ∼50% of the islets initial first responder cells remained consistent during repeated glucose elevation (Fig.S5 A). Therefore, the temporal consistency of this cell state is not rigid. Some of them surrendered their first responder role either to a nearest neighbor cell, or to a 2^nd^ nearest neighbor. Therefore, the spatial location of the first responder cluster remained consistent (Fig.2 E).

We demonstrated that first responder cells are distinct from other functional subpopulations of β-cells previously defined by [Ca^2+^] signatures. Leader (wave-origin) sells have been associated with regulating second phase [Ca^2+^] dynamics [22, 23]. Hub cells have been associated with maintaining elevated, coordinated [Ca^2+^]. Thus, the heterogeneity that controls second-phase [Ca^2+^] is different than the first-phase [Ca^2+^]. We do note that hub cells were previously identified following more rapid measurements of [Ca^2+^] dynamics (∼10fps) than in our study here (∼1fps) [20]. The lack of fast (<1s) timescale [Ca^2+^] dynamics in our analysis may therefore exclude some hub cells and mean that the hub-like (ph2) cells we identify are not exactly analogous to those previously identified.

While the first-responder cells remained in the lead over 1-2 h, this consistency was gradually lost after 24h (Fig.2). The overall responsiveness of the islet was maintained (demonstrated by calcium influx, Fig2. G), indicating the loss of state is not simply due to islet dysfunction. This finding suggests that first responder cells represent a transient functional state of the islet and not a permanent subpopulation of the β-cell. We are not aware of other long-term imaging studies beyond 1-2 h that test whether functional subpopulations represent transient states of the β-cell rather than permanent β-cell subpopulations. The current lack of genetic markers for first responder cells hinders robust lineage tracing approaches to validate this finding. Indeed, this first-responder cell state may represent a sub-set of the more mature functional cell subpopulations that have been genetically marked [2]. Cyclic expression of *Ins2* gene activity has been reported in subsets of β-cells [36], suggesting that transient states of the β-cell can exist. A correlation between *Ins* expression and *GJD2* (coding Cx36) expression has also been demonstrated [37].

Thus, we speculate that fluctuations in Cx36 expression may contribute to first responder cells: increases in Cx36 gap junction conductance would suppress first responder cells as a result of hyper-polarization by less excitable neighboring cells. Decreases in Cx36 gap junction conductance (as is observed in first responder cells, Fig.4) would allow the more excitable first responder cells to respond earlier and impact their neighboring cells. Decreased gap junction conductance would prevent a cell being hyper-polarized and inhibited by less excitable neighboring cells. Our computational model of the islet also showed lower gap junction conductance within the first responder cells. However, we cannot exclude a role for intra-islet paracrine factors in also regulating first responder cells.

An important consideration is that our study, along with other studies [20, 22], examines a single plane of cells within the islet. Thus, the first responder cells are relative to that islet region. It is quite likely that elsewhere in the islet at other planes, cells that respond earlier are present which may play a more important role. Nevertheless, first responder cells represent a distinct functional state of the β-cell, stable for ∼24 hours, and regulating the first phase of [Ca^2+^] and likely also insulin secretion.

### First responder cells represent a more excitable subpopulation

We hypothesized and demonstrated that first responder cells are important to recruit elevated [Ca^2+^] in neighboring cells. Previously we identified highly-excitable cells that effectively recruited elevated [Ca^2+^] in neighboring cells [22]. These cells showed increased NAD(P)H responses. However, while first responder cells showed increased [Ca^2+^] influx compared to neighboring cells, they did not show any difference in NAD(P)H levels (Fig.4). First responder cells however did show some level of consistency following glibeclamide stimulation that inhibits K_ATP_ channels (Fig.3), although not to the consistency found with glucose stimulation. First responder cells likely show differences in ion channel composition, which may include reduced K_ATP_, but equally could include an increased resting depolarizing current (Fig.7A). For example, increased expression of HCN channels has been found in a population of human beta-cells that is important for increased insulin secretion [39]. The reduced correlation between glucose and glibenclamide stimulation suggests that first responders are also defined by other factors. Functional subpopulations (e.g ‘hub cells’, ‘wave-origin’ or ‘leader’) have been identified to show differences in glucose metabolism [20, 22], which may also explain their distinction from first responder cells.

Notably, we observed consistent results within our computational model of the islet (Fig.6). In the model, first responder cells showed decreased K_ATP_ conductance but little difference in GK activity. Supporting this, disrupting K_ATP_ conductance in the first responder cells led to other cells responding earlier. Thus, first responder cells are likely more excitable due to altered ion channel composition, although whether this is due to altered K_ATP_ channel activity or other ion channels (e.g. HCN channels) is still to be determined. We also observed in the model that the cell position within the islet was important: rearranging the position of the first responder cells, and thus the neighboring cells in contact with the first responder cells, disrupted their consistency. While technically challenging, dissociation of islets post-first responder cell identification, and association of islets into pseudo-islet structures would allow this property to be tested [40]. Nevertheless, we did observe that the first-phase response time is spatially organized within 2-3 cell layers. This further suggests spatial positioning, and thus the influence of neighboring cells within this region, is important (Fig.7 C).

As such, first responder cells are more excitable cells, likely as a result of altered ion channel composition, as well as partially decreased gap junction coupling, and/or spatial positioning within the islet.

### First responder cells drive first phase [Ca^2+^] elevation

Following fs laser ablation of first responder cells we observed a decline in the first-phase [Ca^2+^] response, in terms of the level of [Ca^2+^] uptake, the numbers of cells showing elevated [Ca^2+^], the first-phase [Ca^2+^] coordination and the time taken to show elevated [Ca^2+^] (Fig.5), compared to the laser ablation of the non-first responder (control) cell. This indicates that first responder cells are necessary for both recruiting and coordinating elevated [Ca^2+^] following glucose elevation. We observed qualitatively similar results in the islet model after removal of first-responder cells: the integrated [Ca^2+^] elevation over the first phase was diminished (Fig.6). This indicates that gap junction coupling is sufficient to mediate the impact of first responder cells over the islet first-phase [Ca^2+^] response to glucose.

During the second-phase of [Ca^2+^] elevation post-ablation we still observed [Ca^2+^] oscillations. Combining this finding with our discovery that different cells leading different phases of the [Ca^2+^] response to glucose (first responders vs leader), we conclude that first responder cells are more important for recruiting and coordinating [Ca^2+^] during a specific time window, and not for the long-term maintenance of [Ca^2+^] elevation (Fig.7 D). Interestingly, this observation is similar to the regulation of [Ca^2+^] by Cx36 gap junction channels. At elevated glucose almost all β-cells are capable of elevating [Ca^2+^] and insulin release. However, some cells elevate [Ca^2+^] and insulin release more rapidly and some elevate more slowly, likely as a result of being more or less excitable, respectively. As a result, while all cells elevate [Ca^2+^], the differences in their timing prevent a robust first-phase insulin release [16].

Furthermore, the islet responded more slowly upon ablation of first responder cells. This indicated the ability of first responder cells to trigger earlier responses in neighboring cells, not just homogenizing otherwise heterogeneous response times. We therefore suggest that the more excitable first responder cells recruit less excitable cells to elevate [Ca^2+^] in a more coordinated uniform manner, to enhance first-phase [Ca^2+^]. It is worth mentioning that in zebrafish islets, laser ablation of those cells that lead the [Ca^2+^] elevation in response to glucose also disproportionately affected islet-average [Ca^2+^] influx, as compared to laser ablation of control cells [24]. Zebrafish islets in that study were comparable in size to the small mouse islets analyzed in our work. It is possible that these zebrafish “leader” cells are analogous to the first responder cells in mouse islets that we describe in this study.

While model results presented here indicated that gap junction coupling was sufficient for 10% of first responder cells to drive first-phase [Ca^2+^], we do note that more substantial diminishments of [Ca^2+^] influx were observed when greater numbers of earlier responding cells were removed. This suggests the existence of functional redundancy in the ability of earlier responding cells to drive the first-phase [Ca^2+^] elevation. Indeed, following ablation of a first responding cell, the ‘new’ first responding cells were cells that responded earlier than average prior to cell ablation. This further explains how larger islets were relatively resistant to destruction of individual cells. In small islets the much greater impact on [Ca^2+^] likely is a result of both the greater % of the islet that has been disrupted, and that there are fewer earlier responding cells to replace the function of the first responder cell. As noted elsewhere, by restricting our analysis to a single plane, as with other functional subpopulation studies, we are also potentially excluding cells that respond even earlier and would be expected to show greater impact in controlling the first phase [Ca^2+^]. As such future studies should focus on full 3D analysis encompassing the whole islet. Measuring how loss of first responder cells impacts insulin release itself will also be important to establish, particularly to establish whether changes in first-phase [Ca^2+^] impact first-phase insulin release.

### Summary

Several functional β-cell subpopulations have been identified that influence islet function, yet it is currently unknown whether these subpopulations overlap and whether the mechanisms by which they affect the islet function are the same. This understanding has been hindered by a lack of standard procedures in identifying functional subpopulations (e.g. by standard analysis of [Ca^2+^] dynamics, optogenetic stimulation or silencing). Longer-term imaging to track the consistency and state of functional subpopulations has also been missing. We combined high resolution confocal microscopy in islets with β-cell specific [Ca^2+^] sensor expression, together with targeted removal of single cells via 2-photon laser ablation and standardized analysis of [Ca^2+^] coordination. We discovered a distinct functional β-cell state that was stable for ∼24h. The state is characterized by increased electrical excitability and slightly reduced gap junction permeability. This state did not overlap with other previously identified functional subpopulations. We discovered organization of the first-phase [Ca^2+^] response and existence of hierarchy in this response where the second earliest to response cell takes over the role of the first responder cell upon the first responder cell ablation. Removal of first responder cells disproportionately disrupted the response time of the islet and [Ca^2+^] levels during the first phase following glucose elevation, compared to the control cell removal in the size-matched islets. Thus, we have identified a cell state that is functionally important to a first phase of calcium response to glucose in individual islets.

## Methods

### Animal care

Male and female mice were used under protocols approved by the University of Colorado Institutional Animal Care and Use Committee. β-cell-specific GCaMP6s expression (β-GCaMP6s) was achieved through crossing a MIP-CreER (The Jackson Laboratory) and a GCaMP6s line (The Jackson Laboratory). Genotype was verified through qPCR (Transetyx, Memphis, TN). Mice were held in a temperature-controlled environment with a 12 h light/dark cycle and given continuous access to food and water. CreER-mediated recombination was induced by 5 daily doses of tamoxifen (50mg/kg bw in corn oil) delivered IP.

### Islet isolation and culture

Islets were isolated from mice under ketamine/xylazine anesthesia (80 and 16 mg/kg) by collagenase delivery into the pancreas via injection into the bile duct. The collagenase-inflated pancreas was surgically removed and digested. Islets were handpicked and planted into the glass-bottom dishes (MatTek) using CellTak cell tissue adhesive (Sigma-Aldrich). Islets were cultured in RPMI medium (Corning, Tewksbury, MA) containing 10% fetal bovine serum, 100 U/mL penicillin, and 100 mg/mL streptomycin. Islets were incubated at 37C, 5% CO2 for 24-72 h before imaging.

### Imaging

An hour prior to imaging nutrition media from the isolated islets was replaced by an imaging solution (125 mM NaCl, 5.7 mM KCl, 2.5 mM CaCl2, 1.2 mM MgCl2, 10 mM HEPES, and 0.1% BSA, pH 7.4) containing 2 mM glucose. During imaging the glucose level was raised to 11mM. Islets were imaged using either a LSM780 system (Carl Zeiss, Oberkochen, Germany) with a 40x 1.2 NA objective or with an LSM800 system (Carl Zeiss) with 20x 0.8 NA PlanApochromat objective or a 40x 1.2 NA objective, with samples held at 37C.

For [Ca^2+^] measurements GCaMP6s fluorescence was excited using a 488-nm laser. Images were acquired at 1 frame/s at 10-15 um depth from the bottom of the islet. Glucose was elevated 3 minutes after the start of recording, unless stated otherwise.

NAD(P)H autofluorescence and [Ca^2+^] dynamics were performed in the same z-position within the islet. NADH(P)H autofluorescence was imaged under two-photon excitation using a tunable mode-locked Ti:sapphire laser (Chameleon; Coherent, Santa Clara, CA) set to 710 nm. Fluorescence emission was detected at 400–450 nm using the internal detector. Z-stacks of 6–7 images were acquired spanning a depth of 5 μm. First the NAD(P)H was recorded at 2 mM glucose, then the [Ca^2+^] dynamics was recorder at 2 mM and during transition to 11 mM glucose. After the [Ca^2+^] wave was established, the NAD(P)H was recorded at 11 mM glucose.

Cx36 gap junction permeability and [Ca^2+^] dynamics were performed in the same z-position within the islet, with gap junction permeability measured using fluorescence recovery after photobleaching, as previously described [17]. After [Ca^2+^] imaging, islets were loaded with 12 mM Rhodamine-123 for 30 min at 37C in imaging solution. Islets were then washed and FRAP performed at 11mM glucose at room temperature. Rhodamine-123 was excited using a 488-nm laser line, and fluorescence emission was detected at 500–580 nm. Three baseline images were initially recorded. A region of interest was then photobleached achieving, on average, a 50% decrease in fluorescence, and images were then acquired every 5-15 s for 15 min.

### Imaging long term [Ca^2+^] dynamics

The initial [Ca^2+^] dynamics under glucose elevation from 2 to 11 mM at 0 hours was recorded. The dish was marked to indicate its orientation with respect to the microscope stage, and the arrangement of islets was noted to facilitate islet localization in subsequent imaging. After this first time point, imaging solution was replaced by the islet culture media and the dish was kept in the incubator at 37C and 5% CO2 until the next time point. The same cell layer in the islet was imaged at 6h intervals until 48h. For some islets intervals of 12 h were recorded.

### Laser ablation

Laser ablation was performed with two-photon tunable mode-locked Ti:sapphire laser (Chameleon; Coherent, Santa Clara, CA) set to 750 nm. First [Ca^2+^] dynamics was recorded at 2mM and 11 mM glucose, and first responder cells were identified. Then glucose was lowered to 2mM and [Ca^2+^] activity was monitored to ensure the islet returns to a basal level of activity. The first responder cell(s) were identified, and a sub-cell-sized region of interest to be ablated (5×5 μm) was selected either over the first responder cell or over a control cell far from the first responder. Ablation was performed by illuminating the region of interest. [Ca^2+^] dynamics were then imaged during the transition from 2mM to 11mM glucose.

### Analysis of [Ca^2+^] dynamics

We defined first responder cells as 10% of the cells imaged within an islet that responded earliest to show elevated [Ca^2+^]. The time of response was defined as time at which intensity of the fluorescence of the [Ca^2+^] indicator (GCaMP6s) reached half the maximum height following islet stimulation by glucose or other secretagogue. We refer to this time as half-height time, or response time throughout the text.

The leader and lagger cells were defined during the second-phase [Ca^2+^] response to glucose, once oscillations emerge. These cells were defined based on the phase lag of the slow [Ca^2+^] oscillation in each cell with respect to the phase of the average [Ca^2+^] oscillation across the islet, as previously presented [41]. Islets that lacked second-phase [Ca^2+^] oscillations were excluded from the analysis of the oscillatory phase, and only used for first-phase analysis.

For analysis of long term [Ca^2+^] dynamics, only those cells that correspond to cells imaged in the first time point (0 h) were considered. These cells were identified by locating their relative positions with respect to non-β cells which did not express GCaMP6s. Additional cells that appear in the plane of as a result of islets slowly flattening over time were not considered.

### Distances between the cells

X and Y coordinates of centers of each ROI outlining a cell were obtained in Fiji software. Then distances between the cells of interest were calculated using these coordinates. In Fig.1 J distance from first responder to first responder indicates average distance between all first responder cells per islet. For example, in the islet shown in Fig.1 I there were several first responding cells (cell #30, #34, …, #48), the distances were measured from cell#48 to #30, from #30 to #34, etc. And then the average value was plotted as a dot in Fig.1E (red bar). Similarly, distances between each first responder and other cells of interest were calculated and averaged per islet.

### Network analysis

Network connectivity analysis for second-phase [Ca^2+^] response presented in Fig.1 and Fig. S1 was performed by Dr. David Hodson as he reported previously [20] and using the same algorithm. Briefly, [Ca^2+^] time courses were binarized based on intensity deviation from the mean intensity using 20% intensity cutoff. Intensity above this cutoff was assigned a value of 1, and below – value of 0. A co-activity matrix with elements C_ij_ for each cell pair in the islet was constructed from the binarized signal.

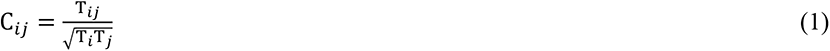

The T_i_ and T_j_ represent time (sec) of activity (when intensity was >20% cutoff) for cells *i* and *j*, and the T_ij_ represents time of co-activity of a cell pair. Then the co-activity matrix was shuffled >9999 times to construct a random co-activity matrix, with elements C_ij_*, which was used to account for a co-activity being due to chance. The experimental co-activity matrix was then adjusted using threshold constructed of the mean value of C_ij_* and a standard deviation from the mean, σ*:

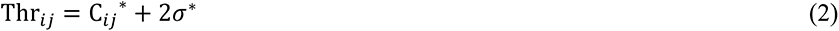

The percent of links was calculated with respect to the maximum number of links per cell in each individual islet. For example, if a most connected cell possessed max=10 links, and other cells had 1, 3, …7 – then % were: 10%, 30%, …70%.

Pearson-product-based network analysis presented in Fig.5 was performed as previously reported [21]. [Ca^2+^] time courses were analyzed during the first-phase [Ca^2+^] response, and the analyzed time ranges were chosen to be equal for pre- and post-ablation. The Pearson product for each cell pair in the islet was calculated over each time point, and the time-average values were computed to construct a correlation matrix. An adjacency matrix was calculated by applying a threshold to the correlation matrix. The same threshold of 0.9 was applied to all islets. All cell pairs with a non-zero values in the adjacency matrix were considered to have a functional link.

### Islet modelling

The coupled β-cell model was described previously [42] and adapted from the published Cha-Noma single cell model [43, 44]. All code was written in C++ and run on the SUMMIT supercomputer (University of Colorado Boulder).

The membrane potential (V_i_) for each β-cell i is related to the sum of individual ion currents as described by [43]:

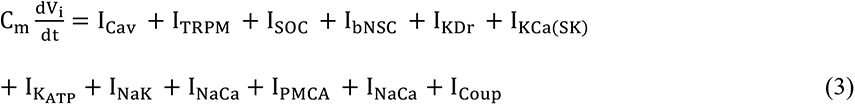

Where the gap junction mediated current I_Coup_ [19] is:

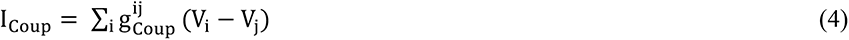

There are N = 1000 cells in each simulation. Heterogeneity was introduced by randomizing multiple variables according to a Gaussian distribution (Supplemental Table). Heterogeneity in Cx36 gap junctions was modeled as a γ-distribution with parameters k=θ=4 as described previously [22] and scaled to an average g_Coup_ between cells = 120 pS.

The flux of glycolysis J_glc_, which is limited by the rate of k_glc_ activity in the β-cell, is described as:

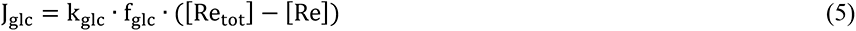

Where k_glc_ is the maximum rate of glycolysis (equivalent to GK activity), which was simulated as a continuous Gaussian distribution with a mean of 0.000126 ms^-1^ and standard deviation of 25% of the mean. [Re_tot_] = 10mM, the total amount of pyrimidine nucleotides. The ATP and glucose dependence of glycolysis (GK activity) is:

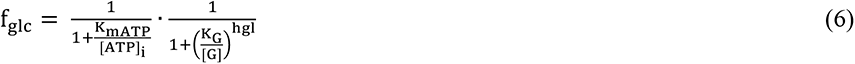

Where [G] is the extracellular concentration of glucose, hgl is the hill coefficient, K_G_ is the half maximal concentration of glucose, and K_mATP_ is the half maximal concentration of ATP.

In simulations where a specific heterogenous parameter is adjusted, the first 100 responding cells were determined in a simulation (see below), and these first responder cells were adjusted so that the parameter of choice (g_KATP_, k_glc_, or g_Coup_) was forced to be at the average value (Supplemental Table). The value of the parameter of the other cells in the islet was adjusted so that the average of the parameter value over the islet remained unchanged. A new simulation with these adjusted parameters was run. As a control, a random set of 100 cells are distributed across the islet, and these random cells were selected with the same spatial organization as first responder cells from a simulation that initiated with a different random number seed. These random cells are adjusted in the same way as described above to the average parameter value, while the other cells in the simulation are adjusted so that the average islet value remains unchanged.

### Islet modelling analysis

All simulation data analysis was performed using custom MATLAB scripts. First responder cells were determined using the [Ca^2+^] time courses of each cell during the first 20-72 seconds, which covers the first phase [Ca^2+^] response. The response time was calculated as the time to half the maximum [Ca^2+^] level. First responder cells were then determined to be the 100 (10%) cells with the earliest response time.

In simulations where cells are uncoupled, a given % of cells with the lowest response time are uncoupled from the simulation, but the next 100 cells are analyzed as the first responders of this new simulation. To uncouple cells from the simulated islet, the conductance, g_Coup_, of the cells to be removed is set to 0 pS. Removed cells are excluded from subsequent islet analysis. For control simulations a random % of cells are uncoupled from the islet. These cells are distributed across the islet in the same organization as the first responder cells from a different simulation.

### Statistical analysis

All statistical analysis was performed in Prism (GraphPad). For computational results a one-way repeated measures ANOVA with Tukey post-hoc analysis was utilized to test for significant differences between either the WT simulation or a ‘Random’ control simulation that matched islets before and after parameters are adjusted or cells are uncoupled. For experimental results either one-way ANOVA, or a two-tailed t-test were used (indicated in figure captions) to compare parameters of a specific β-cell subpopulation to the corresponding islet-average parameters. Data is reported as mean ± s.e.m. (standard error in the mean) unless otherwise indicated. Linear regression analysis was presented as the trend with 95% confidence intervals. The difference of the slope from zero was evaluated with the F test and the p value was reported.

### Sample size estimate

Power analysis was performed to estimate sample size required for each set of experiment. For example, for experimental set of the relative location of subpopulations (Fig.1D) mean and standard deviation of the distribution of distances from first responder cell to every other first responder in the same islet was chosen as the “expected” group. Power analysis yielded the n=6 per group required to provide the effect size of 20% assuming symmetrical distribution. For the % of functional links (Fig.1) mean and standard deviation of the islet-median % of links distribution was chosen as the “expected” group. Power analysis yielded the n=8 per group required to provide the effect size of 20% assuming symmetrical distribution. In the same fashion mean and standard deviation of the pilot “expected” group were used for power analysis of the rest of the experimental sets (short- and long-term consistency of the response time of the 1^st^ responder, difference in the electrical coupling measured with FRAP, effect of the laser ablation etc). The probability of the type1 error, alpha was chosen as 5%, power was chosen as 80%, and the effect size of 20% was used to estimate sample size through the power analysis.

## Acknowledgments

Richard KP Benninger (University of Colorado) is the guarantor of this work and, as such, had full access to all the data in the study and takes responsibility for the integrity of the data and the accuracy of the data analysis. All authors acknowledge that no conflict of interest exists. This work was supported by Juvenile Diabetes Research Foundation (JDRF) Grant 5-CDA-2014-198-A-N; National Institute of Health (NIH) grants R01 DK102950, R01 DK106412 (to RKPB); by JDRF grant 3-PDF-2019-741-A-N (to VK); by NIH grant F31 DK126360 (to JMD). D.J.H. was supported by a Diabetes UK R.D. Lawrence (12/0004431) Fellowship, a Welcome Trust Institutional Support Award, and MRC (MR/N00275X/1 and MR/S025618/1) and Diabetes UK (17/0005681) Project Grants. This project has received funding from the European Research Council (ERC) under the European Union’s Horizon 2020 research and innovation program (Starting Grant 715884 to D.J.H.). The funders had no role in the study design, data collection and analysis, decisions to publish, or preparation of the manuscript. We appreciate helpful information on Pearson-based network analysis form Dr. Andraz Stozer, Institute of Physiology, University of Maribor.

## Author Contributions

VK conceived of the idea, designed and performed experiments and computer simulations, analyzed data, wrote the manuscript; JMD performed computer simulations, analyzed data, edited text; WES performed animal surgeries, analyzed data; DJH analyzed data, edited the manuscript; RAP performed animal surgeries; AD analyzed data; MSI analyzed data; RKPB conceived of the idea, designed experiments, and wrote the manuscript.

## List of Source Data Files

Figure 1 – Source Data (values used for all graphs in Figure 1)

Figure 2 – Source Data (values used for all graphs in Figure 2)

Figure 3 – Source Data (values used for all graphs in Figure 3)

Figure 4 – Source Data (values used for all graphs in Figure 4)

Figure 5 – Source Data (values used for all graphs in Figure 5)

Figure 6 – Source Data (values used for all graphs in Figure 6)

Statistical analysis (Summary of statistical analysis for Figures 1-6)

KATP_MovetoAvg_seed1 (model code for Figure 6 I, J)

Coup_MovetoAvg_seed1 (model code for Figure 6 I, J)

Uncouple_FirstResp_Seed1_perc20 (model code for Figure 6 L)

---

**Figure S1.**
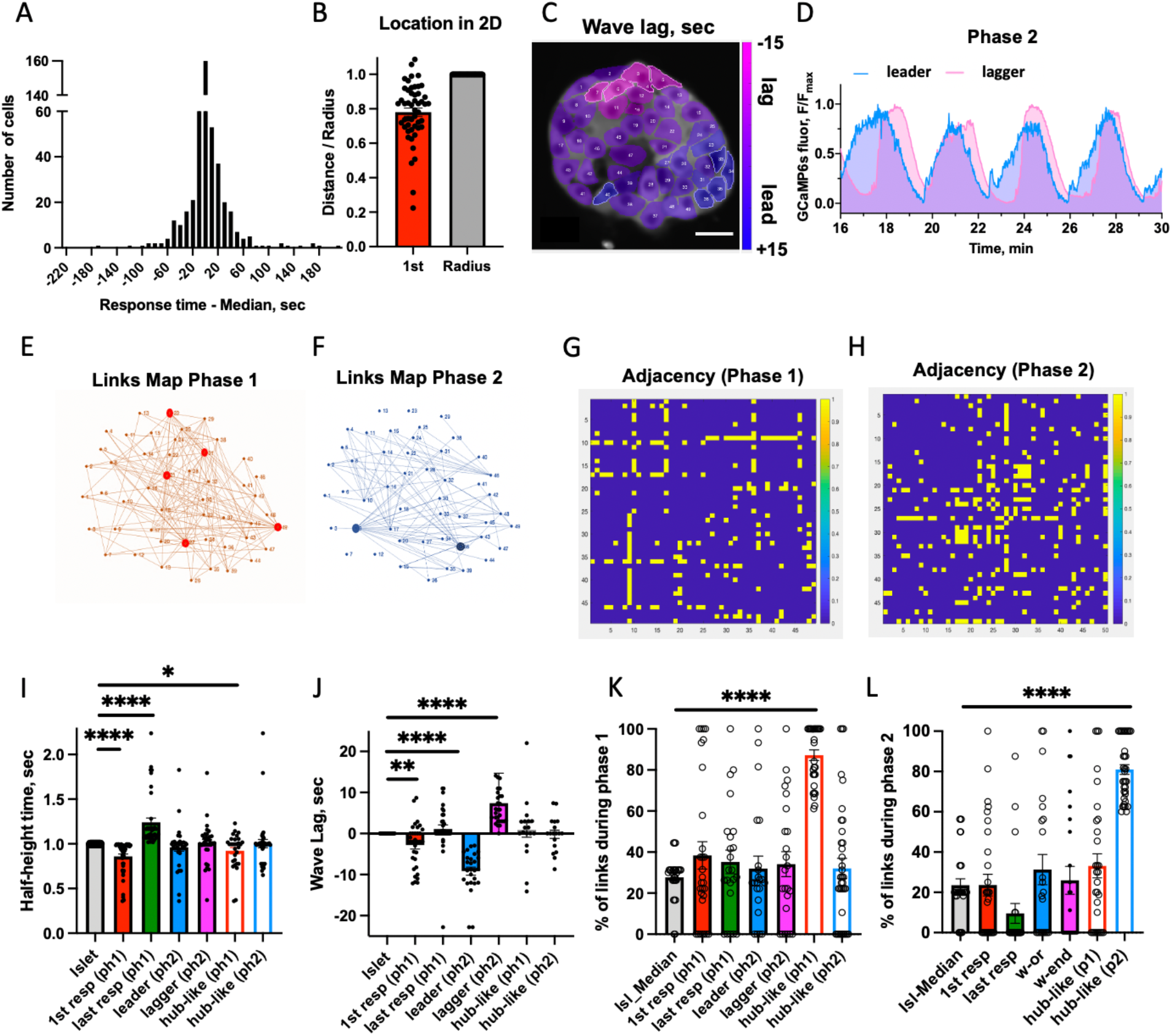
A) Cumulative response time (Tresp) distribution for 19 islets. B) Location of 1^st^ responder cells in the 2D islet plane, normalized by islet radius. C) Representative false-color map of calcium phase lag. D) Example of calcium leader and “lagger” cell time traces. E) and F) link maps obtained using network analysis for first- and second-phase calcium response, correspondingly. G) and H) Adjacency matrices for for first- and second-phase calcium response, correspondingly. I) [Ca^2+^] response time to glucose elevation for different beta-cell states (n=8-10 islets, m=30-45 cells). J) Phase lag of the Ca^2+^ wave with respect to the islet-average wave for different beta-cell states (n=8-12 islets, m=18-34 cells). K), L) Coordination (network) analysis for n=8 islets performed for first- and second-phase calcium dynamics, correspondingly. Functional network analysis was performed via binarization and co-activity matrix analysis, as described before [20]. Statistical tests: I, J: one-sample t-test, K, L: ordinary one-way ANOVA where **** represents p<0.0001, *** p<0.0002, ** p<0.0021, * p<0.0332 indicated for comparison of the groups.

**Figure S2.**
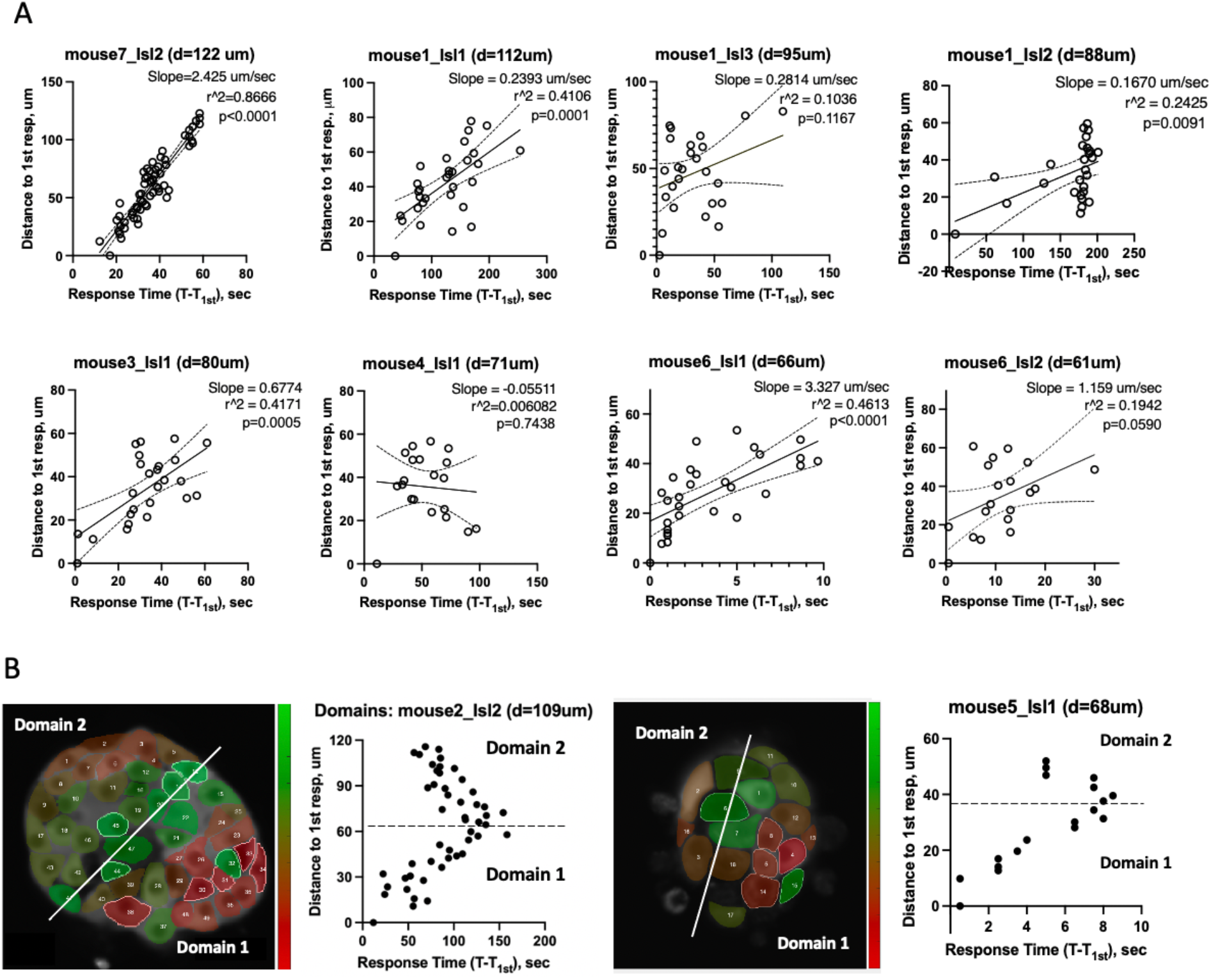
A) Correlation between the absolute response time of each beta-cell in an islet and their proximity to the first responder beta-cell, for 8 islets. Solid line indicates regression, dashed line indicates 95% CI. E) Examples of islets in which there are 2 domains with local first responder clusters per each domain, as well as distance of each cell in the islet plane to 1st responders (located in Domain1) vs time of response to glucose.

**Figure S3.**
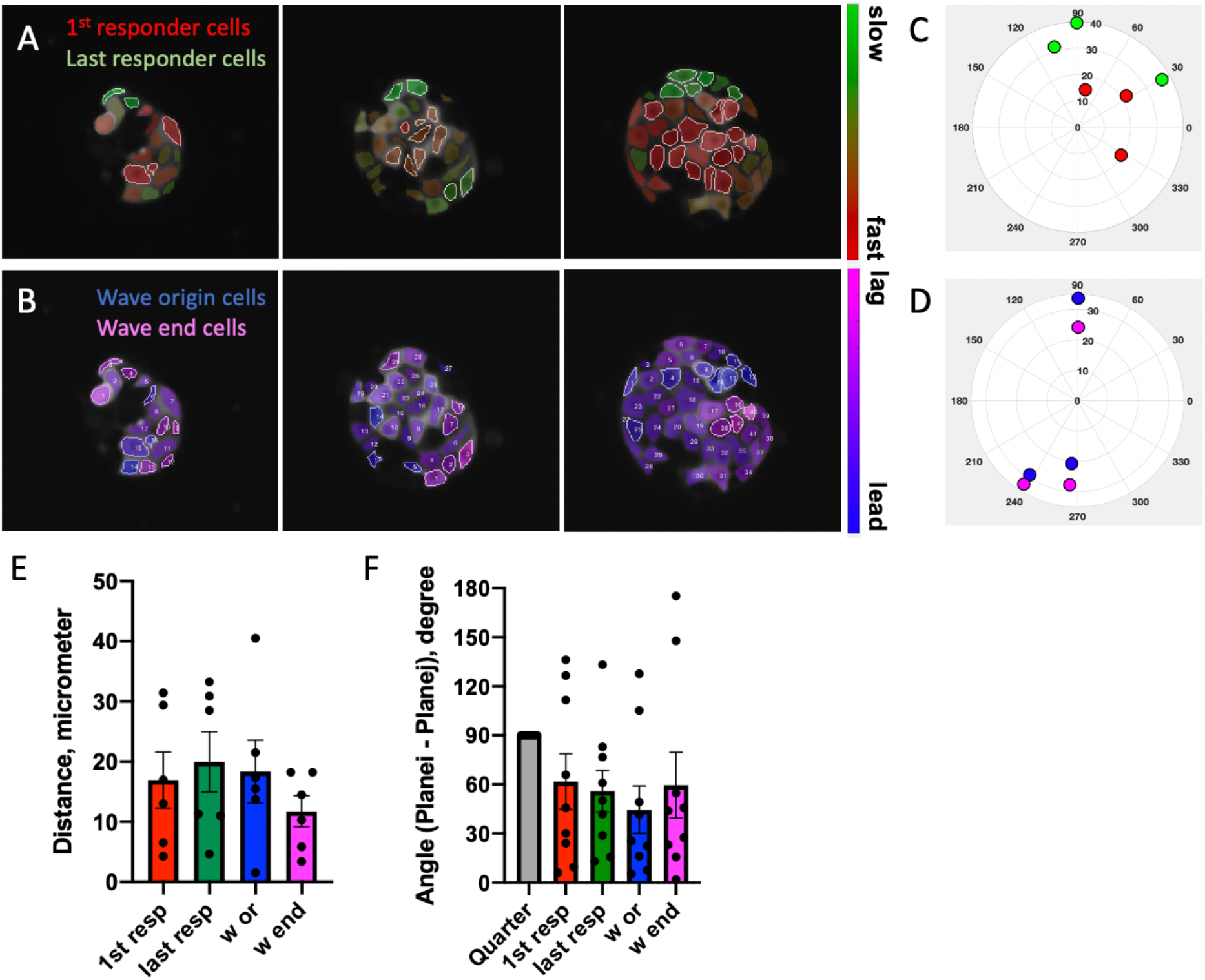
A) Pseudo-color map representing time of response to glucose during the first-phase [Ca^2+^] response in three adjacent planes in the islet separated by 10 μm. First and last responder cells are highlighted with the white borders. B) Pseudo-color map representing wave propagation in the same islet planes during the second-phase [Ca^2+^] response. Wave origin and wave end cells are highlighted with the white borders. C), D) Plane-average positions of each beta cell subpopulation for three planes. E) Distances between the plane-average positions of each subpopulation (position in plane (i) – position in plane (j)). Average distances are ∼20 μm, indicating that location of all four subpopulations of interest is conserved in 3D. F) Polar angle between the plane-average positions of the subpopulations (angle in plane (i) – angle in plane (j)). Average angles are below 90 degrees, indicating spatial conservation of the location of all four subpopulations of interest in 3D. For (E,F) n=4 islets were studied.

**Figure S4.**
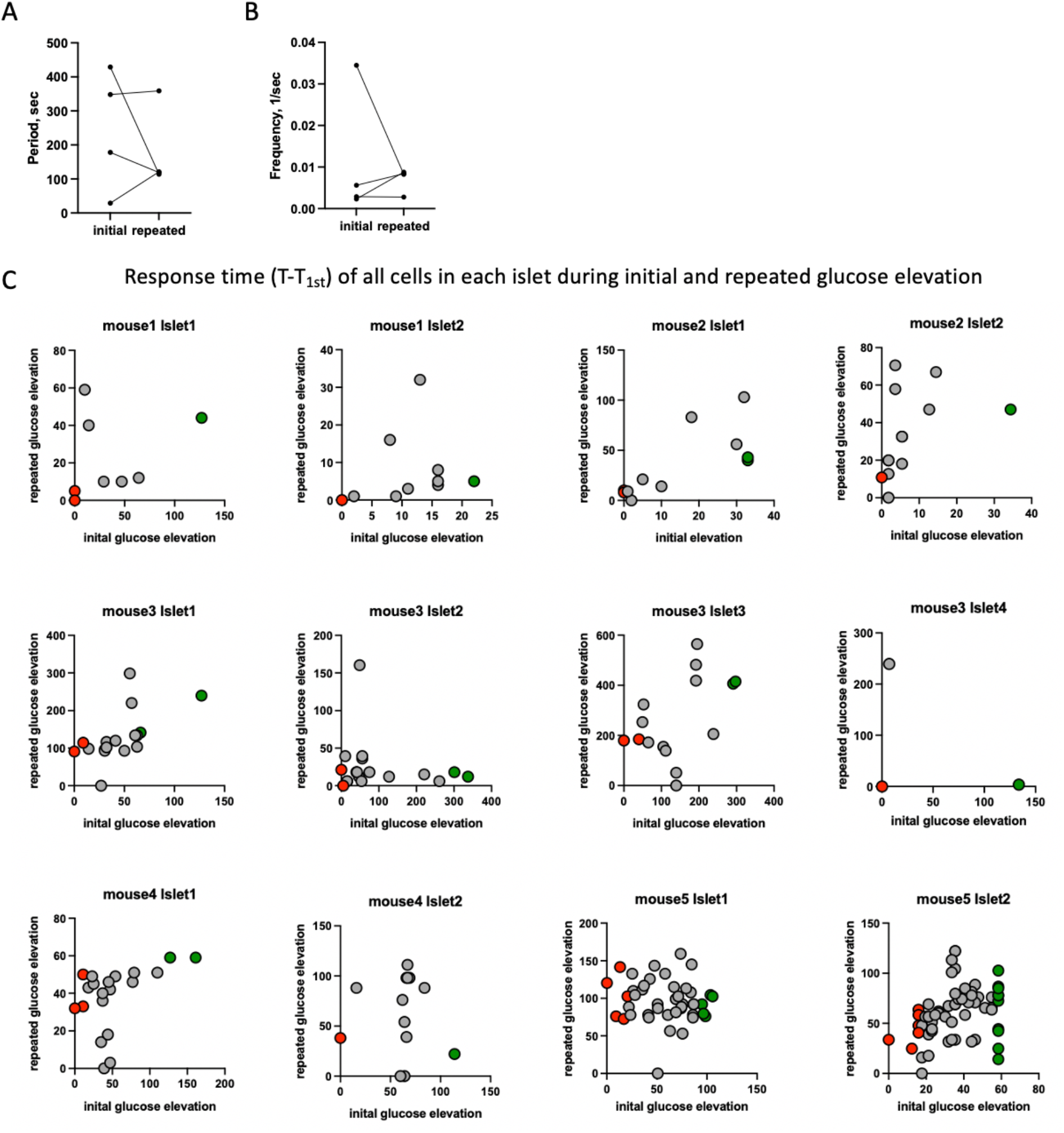
A) Quantification of the Ca^2+^ oscillation period and B) frequency during initial and repeated glucose elevation (n=4 islets). No significant difference was found in one sample t-test with initial-repeated period (frequency) difference compared to 0. C) Response time of all cells in the optical section of the islet during initial vs during repeated glucose elevation. Red dots represent first responder, and green – last responder cells identified during the initial glucose elevation.

**Figure S5.**
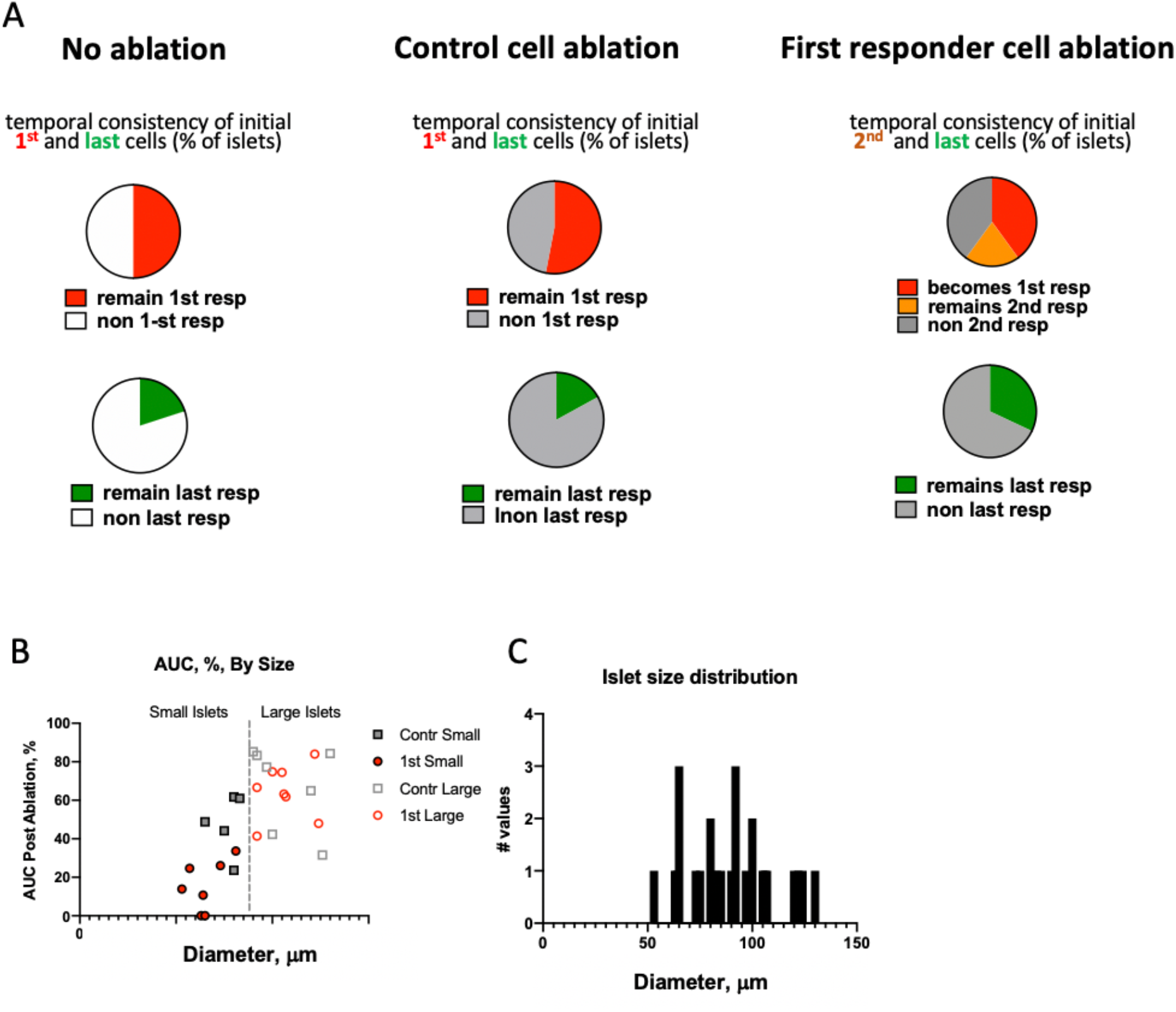
A) Left Pie chart: Percent of islets in which initial first and last responders remain in their role during repeated glucose elevation. Middle Pie chart: same as Left but following control cell ablation. Right Pie charts: same as middle but following first responder cell ablation. B) Islet size dependence of the [Ca^2+^] influx into the islet (area under the curve of the [Ca^2+^] time-course) for random and first responder ablation cases. D) Size distribution of the islets studied in the ablation experiments.

**Supplemental Table.**
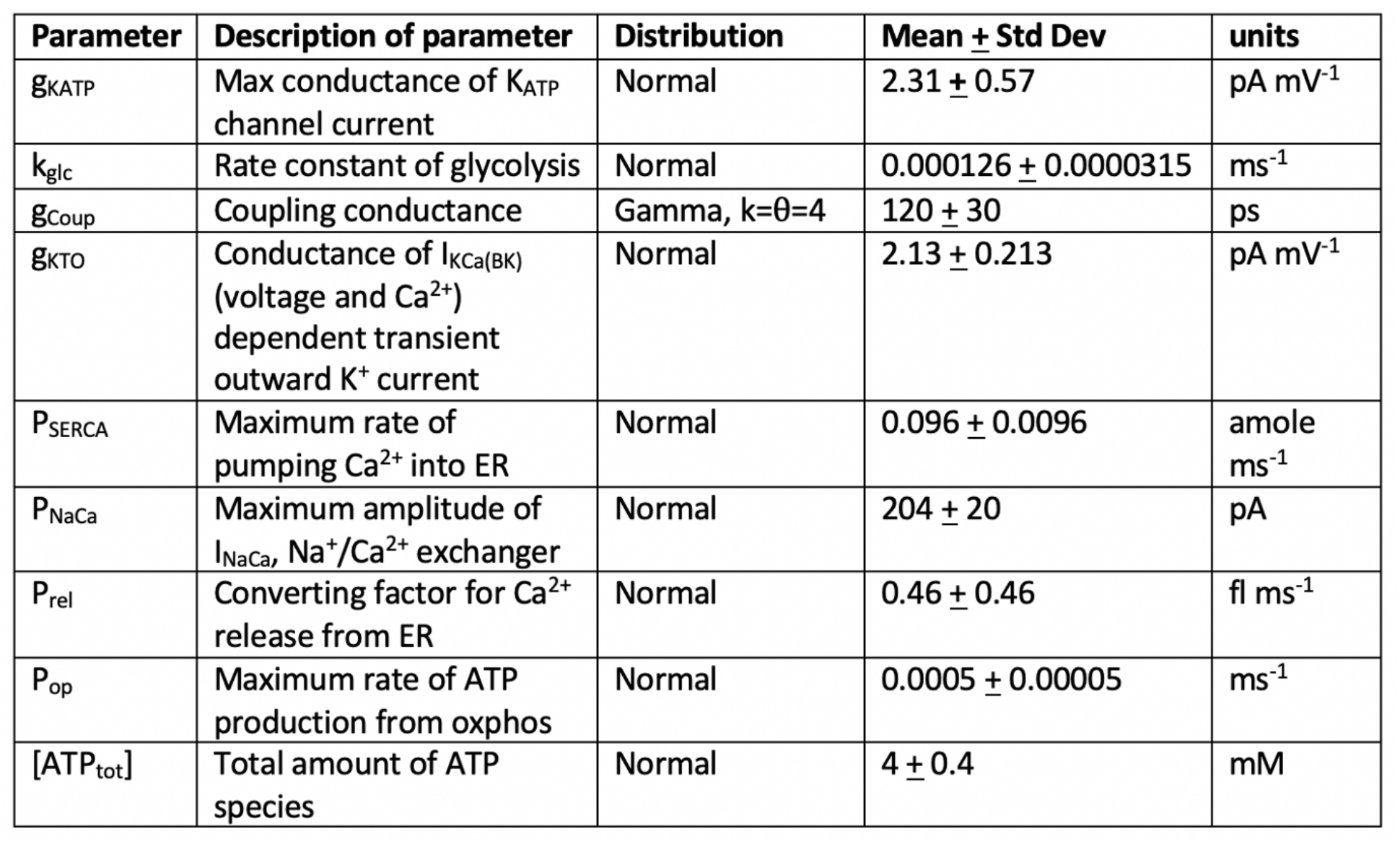
Values of heterogeneous parameters used in the islet simulations.

| Parameter | Description of parameter | Distribution | Mean ± Std Dev | units |
| --- | --- | --- | --- | --- |
| $g_{KATP}$ | Max conductance of $K_{ATP}$ channel current | Normal | $2.31 \pm 0.57$ | $\text{pA mV}^{-1}$ |
| $k_{glc}$ | Rate constant of glycolysis | Normal | $0.000126 \pm 0.0000315$ | $\text{ms}^{-1}$ |
| $g_{Coup}$ | Coupling conductance | Gamma, $k=\theta=4$ | $120 \pm 30$ | ps |
| $g_{KTO}$ | Conductance of $I_{KCa(BK)}$ (voltage and $\text{Ca}^{2+}$) dependent transient outward $\text{K}^+$ current | Normal | $2.13 \pm 0.213$ | $\text{pA mV}^{-1}$ |
| $P_{SERCA}$ | Maximum rate of pumping $\text{Ca}^{2+}$ into ER | Normal | $0.096 \pm 0.0096$ | $\text{amole ms}^{-1}$ |
| $P_{NaCa}$ | Maximum amplitude of $I_{NaCa}$, $\text{Na}^+/\text{Ca}^{2+}$ exchanger | Normal | $204 \pm 20$ | pA |
| $P_{rel}$ | Converting factor for $\text{Ca}^{2+}$ release from ER | Normal | $0.46 \pm 0.46$ | $\text{fl ms}^{-1}$ |
| $P_{op}$ | Maximum rate of ATP production from oxphos | Normal | $0.0005 \pm 0.00005$ | $\text{ms}^{-1}$ |
| $[\text{ATP}_{tot}]$ | Total amount of ATP species | Normal | $4 \pm 0.4$ | mM |

## References

1. D. Salomon and P. Meda, “Heterogeneity and contact-dependent regulation of hormone secretion by individual B cells,” (in eng), Exp Cell Res, vol. 162, no. 2, pp. 507–20, Feb 1986, doi: 10.1016/0014-4827(86)90354-x.

2. E. Bader et al., “Identification of proliferative and mature β-cells in the islets of Langerhans,” (in eng), Nature, vol. 535, no. 7612, pp. 430–4, 07 2016, doi: 10.1038/nature18624.

3. H. Katsuta et al., “Subpopulations of GFP-marked mouse pancreatic β-cells differ in size, granularity, and insulin secretion,” (in eng), Endocrinology, vol. 153, no. 11, pp. 5180-7, Nov 2012, doi: 10.1210/en.2012-1257.

4. M. Karaca et al., “Exploring functional beta-cell heterogeneity in vivo using PSA-NCAM as a specific marker,” (in eng), PLoS One, vol. 4, no. 5, p. e5555, 2009, doi: 10.1371/journal.pone.0005555.

5. Å. Segerstolpe et al., “Single-Cell Transcriptome Profiling of Human Pancreatic Islets in Health and Type 2 Diabetes,” (in eng), Cell Metab, vol. 24, no. 4, pp. 593-607, 10 2016, doi: 10.1016/j.cmet.2016.08.020.

6. Y. J. Wang et al., “Multiplexed In Situ Imaging Mass Cytometry Analysis of the Human Endocrine Pancreas and Immune System in Type 1 Diabetes,” (in eng), Cell Metab, vol. 29, no. 3, pp. 769–783.e4, 03 2019, doi: 10.1016/j.cmet.2019.01.003.

7. L. Michon et al., “Involvement of gap junctional communication in secretion,” (in eng), Biochim Biophys Acta, vol. 1719, no. 1-2, pp. 82–101, Dec 2005, doi: 10.1016/j.bbamem.2005.11.003.

8. S. Bavamian et al., “Islet-cell-to-cell communication as basis for normal insulin secretion,” (in eng), Diabetes Obes Metab, vol. 9 Suppl 2, pp. 118-32, Nov 2007, doi: 10.1111/j.1463-1326.2007.00780.x.

9. V. Serre-Beinier et al., “Cx36 preferentially connects beta-cells within pancreatic islets,” (in eng), Diabetes, vol. 49, no. 5, pp. 727–34, May 2000, doi: 10.2337/diabetes.49.5.727.

10. R. K. Benninger, M. Zhang, W. S. Head, L. S. Satin, and D. W. Piston, “Gap junction coupling and calcium waves in the pancreatic islet,” (in eng), Biophys J, vol. 95, no. 11, pp. 5048–61, Dec 2008, doi: 10.1529/biophysj.108.140863.

11. M. A. Ravier et al., “Loss of connexin36 channels alters beta-cell coupling, islet synchronization of glucose-induced Ca2+ and insulin oscillations, and basal insulin release,” (in eng), Diabetes, vol. 54, no. 6, pp. 1798–807, Jun 2005, doi: 10.2337/diabetes.54.6.1798.

12. A. P. Moreno, V. M. Berthoud, G. Pérez-Palacios, and E. M. Pérez-Armendariz, “Biophysical evidence that connexin-36 forms functional gap junction channels between pancreatic mouse beta-cells,” (in eng), Am J Physiol Endocrinol Metab, vol. 288, no. 5, pp. E948–56, May 2005, doi: 10.1152/ajpendo.00216.2004.

13. S. Speier, A. Gjinovci, A. Charollais, P. Meda, and M. Rupnik, “Cx36-mediated coupling reduces beta-cell heterogeneity, confines the stimulating glucose concentration range, and affects insulin release kinetics,” (in eng), Diabetes, vol. 56, no. 4, pp. 1078-86, Apr 2007, doi: 10.2337/db06-0232.

14. R. K. Benninger, W. S. Head, M. Zhang, L. S. Satin, and D. W. Piston, “Gap junctions and other mechanisms of cell-cell communication regulate basal insulin secretion in the pancreatic islet,” (in eng), J Physiol, vol. 589, no. Pt 22, pp. 5453–66, Nov 2011, doi: 10.1113/jphysiol.2011.218909.

15. J. V. Rocheleau et al., “Critical role of gap junction coupled KATP channel activity for regulated insulin secretion,” (in eng), PLoS Biol, vol. 4, no. 2, p. e26, Feb 2006, doi: 10.1371/journal.pbio.0040026.

16. W. S. Head, M. L. Orseth, C. S. Nunemaker, L. S. Satin, D. W. Piston, and R. K. Benninger, “Connexin-36 gap junctions regulate in vivo first- and second-phase insulin secretion dynamics and glucose tolerance in the conscious mouse,” (in eng), Diabetes, vol. 61, no. 7, pp. 1700–7, Jul 2012, doi: 10.2337/db11-1312.

17. N. L. Farnsworth, A. Hemmati, M. Pozzoli, and R. K. Benninger, “Fluorescence recovery after photobleaching reveals regulation and distribution of connexin36 gap junction coupling within mouse islets of Langerhans,” (in eng), J Physiol, vol. 592, no. 20, pp. 4431–46, Oct 2014, doi: 10.1113/jphysiol.2014.276733.

18. D. G. Pipeleers, “Heterogeneity in pancreatic beta-cell population,” (in eng), Diabetes, vol. 41, no. 7, pp. 777–81, Jul 1992, doi: 10.2337/diab.41.7.777.

19. T. H. Hraha, M. J. Westacott, M. Pozzoli, A. M. Notary, P. M. McClatchey, and R. K. Benninger, “Phase transitions in the multi-cellular regulatory behavior of pancreatic islet excitability,” PLoS Comput Biol, vol. 10, no. 9, p. e1003819, Sep 2014, doi: 10.1371/journal.pcbi.1003819.

20. N. R. Johnston et al., “Beta Cell Hubs Dictate Pancreatic Islet Responses to Glucose,” (in eng), Cell Metab, vol. 24, no. 3, pp. 389–401, 09 2016, doi: 10.1016/j.cmet.2016.06.020.

21. A. Stožer et al., “Functional connectivity in islets of Langerhans from mouse pancreas tissue slices,” (in eng), PLoS Comput Biol, vol. 9, no. 2, p. e1002923, 2013, doi: 10.1371/journal.pcbi.1002923.

22. M. J. Westacott, N. W. F. Ludin, and R. K. P. Benninger, “Spatially Organized beta-Cell Subpopulations Control Electrical Dynamics across Islets of Langerhans,” Biophys J, vol. 113, no. 5, pp. 1093–1108, Sep 5 2017, doi: 10.1016/j.bpj.2017.07.021.

23. R. K. Benninger et al., “Intrinsic islet heterogeneity and gap junction coupling determine spatiotemporal Ca²⁺ wave dynamics,” (in eng), Biophys J, vol. 107, no. 11, pp. 2723–33, Dec 2014, doi: 10.1016/j.bpj.2014.10.048.

24. V. Salem et al., “Leader β-cells coordinate Ca,” (in eng), Nat Metab, vol. 1, no. 6, pp. 615–629, Jun 2019, doi: 10.1038/s42255-019-0075-2.

25. D. L. Curry, L. L. Bennett, and G. M. Grodsky, “Dynamics of insulin secretion by the perfused rat pancreas,” (in eng), Endocrinology, vol. 83, no. 3, pp. 572–84, Sep 1968, doi: 10.1210/endo-83-3-572.

26. D. Porte and A. A. Pupo, “Insulin responses to glucose: evidence for a two pool system in man,” (in eng), J Clin Invest, vol. 48, no. 12, pp. 2309–19, Dec 1969, doi: 10.1172/JCI106197.

27. J. C. Henquin, “Relative importance of extracellular and intracellular calcium for the two phases of glucose-stimulated insulin release: studies with theophylline,” (in eng), Endocrinology, vol. 102, no. 3, pp. 723–30, Mar 1978, doi: 10.1210/endo-102-3-723.

28. C. S. Nunemaker, D. H. Wasserman, O. P. McGuinness, I. R. Sweet, J. C. Teague, and L. S. Satin, “Insulin secretion in the conscious mouse is biphasic and pulsatile,” (in eng), Am J Physiol Endocrinol Metab, vol. 290, no. 3, pp. E523-9, Mar 2006, doi: 10.1152/ajpendo.00392.2005.

29. C. S. Nunemaker et al., “Individual mice can be distinguished by the period of their islet calcium oscillations: is there an intrinsic islet period that is imprinted in vivo?,” (in eng), Diabetes, vol. 54, no. 12, pp. 3517–22, Dec 2005, doi: 10.2337/diabetes.54.12.3517.

30. C. S. Nunemaker et al., “Glucose metabolism, islet architecture, and genetic homogeneity in imprinting of [Ca2+](i) and insulin rhythms in mouse islets,” (in eng), PLoS One, vol. 4, no. 12, p. e8428, Dec 2009, doi: 10.1371/journal.pone.0008428.

31. M. Zhang, P. Goforth, R. Bertram, A. Sherman, and L. Satin, “The Ca2+ dynamics of isolated mouse beta-cells and islets: implications for mathematical models,” (in eng), Biophys J, vol. 84, no. 5, pp. 2852–70, May 2003, doi: 10.1016/S0006-3495(03)70014-9.

32. J. Zmazek et al., “Assessing Different Temporal Scales of Calcium Dynamics in Networks of Beta Cell Populations,” (in eng), Front Physiol, vol. 12, p. 612233, 2021, doi: 10.3389/fphys.2021.612233.

33. A. Vogel and V. Venugopalan, “Mechanisms of pulsed laser ablation of biological tissues,” (in eng), Chem Rev, vol. 103, no. 2, pp. 577–644, Feb 2003, doi: 10.1021/cr010379n.

34. J. M. Dwulet et al., “How Heterogeneity in Glucokinase and Gap-Junction Coupling Determines the Islet [Ca,” (in eng), Biophys J, vol. 117, no. 11, pp. 2188–2203, 12 2019, doi: 10.1016/j.bpj.2019.10.037.

35. R. K. P. Benninger and D. J. Hodson, “New Understanding of β-Cell Heterogeneity and In Situ Islet Function,” (in eng), Diabetes, vol. 67, no. 4, pp. 537–547, 04 2018, doi: 10.2337/dbi17-0040.

36. H. Modi, “Ins2 gene bursting activity defines a mature β-cell state,” BioRxiv, 2019.

37. V. Serre-Beinier et al., “Cx36 makes channels coupling human pancreatic beta-cells, and correlates with insulin expression,” (in eng), Hum Mol Genet, vol. 18, no. 3, pp. 428–39, Feb 2009, doi: 10.1093/hmg/ddn370.

38. J. M. Dwulet, J. K. Briggs, and R. K. P. Benninger, “Small subpopulations of β-cells do not drive islet oscillatory [Ca 2+] dynamics via gap junction communication,” vol. https://doi.org/10.1101/2020.10.28.358457, ed. BioRXiv, 2020.

39. C. Dorrell et al., “Human islets contain four distinct subtypes of β cells,” (in eng), Nat Commun, vol. 7, p. 11756, 07 2016, doi: 10.1038/ncomms11756.

40. K. Shekiro, T. H. Hraha, A. B. Bernard, R. K. Benninger, and K. S. Anseth “Engineering Functional Pseudo-Islets of Defined Sizes from Primary Murine Cells Using PEG Microwell Devices,” ed, 2020.

41. M. J. Westacott, N. W. F. Ludin, and R. K. P. Benninger, “Spatially Organized β-Cell Subpopulations Control Electrical Dynamics across Islets of Langerhans,” (in eng), Biophys J, vol. 113, no. 5, pp. 1093–1108, Sep 2017, doi: 10.1016/j.bpj.2017.07.021.

42. J. M. Dwulet et al., “How Heterogeneity in Glucokinase and Gap-Junction Coupling Determines the Islet [Ca(2+)] Response,” Biophys J, vol. 117, no. 11, pp. 2188–2203, Dec 3 2019, doi: 10.1016/j.bpj.2019.10.037.

43. C. Y. Cha et al., “Ionic mechanisms and Ca2+ dynamics underlying the glucose response of pancreatic beta cells: a simulation study,” (in English), J Gen Physiol, vol. 138, no. 1, pp. 21-37, Jul 2011, doi: DOI 10.1085/jgp.201110611.

44. C. Y. Cha, E. Santos, A. Amano, T. Shimayoshi, and A. Noma, “Time-dependent changes in membrane excitability during glucose-induced bursting activity in pancreatic beta cells,” (in eng), J Gen Physiol, vol. 138, no. 1, pp. 39–47, Jul 2011, doi: 10.1085/jgp.201110612.

